# Modeling Resistance and Recurrence Patterns of Combined Targeted-Chemoradiotherapy Predicts Benefit of Shorter Induction Period

**DOI:** 10.1101/861518

**Authors:** David M. McClatchy, Henning Willers, Aaron N. Hata, Zofia Piotrowska, Lecia V Sequist, Harald Paganetti, Clemens Grassberger

## Abstract

The optimal integration of molecularly targeted therapies with concurrent chemotherapy and radiation (CRT) to improve cures in genotype-defined cancers is unknown. Here, we develop a bio-mathematical framework to simulate patterns of local versus distant recurrences in a heterogeneous lung cancer population receiving combined targeted therapy and CRT. Our validated model predicts that targeted induction before CRT, an approach currently being tested in clinical trials, may render adjuvant targeted therapy less effective due to proliferation of drug-resistant cancer cells when using very long induction periods. Furthermore, our simulations not only demonstrate the competing effects of drug-resistant cell expansion versus overall tumor regression as a function of induction length, but can also directly estimate the probability of observing an improvement in progression-free survival at a given cohort size. We thus demonstrate that such stochastic biological simulations have the potential to quantitatively inform the design of multimodality clinical trials in genotype-defined cancers.

## Introduction

Therapies targeting specific oncogenic driver mutations have dramatically improved survival in patients with metastatic cancer across several disease sites, most significantly in non-small cell lung cancer (NSCLC) harboring mutations in the epidermal growth factor receptor (EGFR) (1, 2). Unfortunately, disease inevitably recurs as pre-existing and *de-novo* drug resistant cells outcompete drug sensitive cells, manifesting the evolution of drug resistant tumors throughout the course of treatment (3-6). In contrast to metastatic EGFR-mutant NSCLC, the role of EGFR inhibitors in locally advanced (LA), non-metastatic NSCLC remains unknown. The addition of EGFR tyrosine kinase inhibitors (TKIs) to standard concurrent chemotherapy and radiation (CRT) in patients with EGFR-mutant cancers has the potential to improve long term survival in this cohort. The recent PACIFIC trial has demonstrated the dramatic benefit that integration of new systemic agents can yield in LA-NSCLC (18): using adjuvant durvalumab, an anti-PD-L1 checkpoint immunotherapeutic agent, and standard-of-care CRT improved progression-free and overall survival significantly. However, this benefit was less pronounced in the subset of EGFR mutant patients (19, 20). Therefore, the PACIFIC trial exemplifies not only the possible benefits of including successful systemic agents into earlier stages of NSCLC, but also the considerable but underutilized potential TKIs may play in the advancement of treatment options for EGFR-mutant LA-NSCLC.

However, it is entirely unknown how to optimally administer TKIs with CRT in order to minimize the risk of acquired drug resistance and improve the efficacy of CRT. Consequently, the design of these multimodality therapy trials is largely empirical with great variability in treatment protocols and a “one-size fits all” approach. Additionally, a large part of the treatment design space is not explored due to the infeasibility of running randomized controlled trials for every possible combination of treatments.

One useful method to sample treatment design space is bio-mathematical modeling, which enables the quantitation of abstract, interconnected phenomena making it a powerful tool in the field of translational oncology for both hypothesis testing and generation (7, 8). Specifically, mathematical modeling of tumor evolution has significantly impacted our understanding and interpretation of acquired resistance to targeted cancer therapies (9-11) and has helped formulate the idea of collateral sensitives to sequential drug regimens (12, 13). Additionally, combing evolutionary modeling of tumors with mechanistic biological models of radiation cell kill and plasma level drug concentrations have enabled novel dosing schedules of radiation therapy in glioblastoma (14), intercalated administration of multiple targeted and chemotherapy agents for melanoma (10), and pulsed injections of targeted therapy in lung cancer (15). Recently, a drug delivery protocol designed using mathematical modeling has been translated into a clinical trial (16).

In this study, we present a generalized bio-mathematical model to optimize TKI plus CRT multimodal regimens in EGFR-mutant LA-NSCLC, with a focus on the relative effect of therapy on local versus occult distant disease sites. In metastatic EGFR mutant NSCLC populations, TKIs result in significant improvements in overall response and survival compared to chemotherapy, but acquired drug resistance inevitable leads to disease recurrence (1, 2, 5, 6, 17). Optimal combination regimens with EGFR TKIs and CRT for LA-NSCLC patients have not yet been studied. A major complicating factor is that failure after CRT can be local or distant, which may be differentially impacted by TKIs. The goal of this study is to inform the design TKI-CRT multimodal clinical trials through mechanistic, bio-mathematical modeling, which encompasses fundamental principles of evolutionary targeted drug resistance, radiation biology, and comorbidities. The model aims at quantitative, *in-silico* analyses of varying treatment design parameters to predict effect sizes in heterogeneous patient populations, and also the probability of observing such effects at a given sample size. We achieve this by first calibrating our model using several institutional datasets, and subsequently demonstrate accurate model predictions of recurrence rates in independent validation datasets of recent multicenter clinical trials and meta-analyses. Next, we exhaustively explore the multimodal treatment design space and show that a TKI induction period of 2-3 months, as used in the recent clinical trials NCT01553942 and NCT01822496 may reduce the effectiveness of adjuvant TKI therapy due to the proliferation of TKI resistant clones during the induction period. Instead, we propose and provide a quantitative rationale for an individualized induction period tailored to resistance evolution. This mathematical framework can not only inform the design of future multimodal TKI-CRT trials, but also be extended to other oncogene driven cancers to design more precise and personalized cancer treatment regimens.

## Results

### Local and Distant NSCLC Progression Model

Our tumor progression model is an advancement of our previously published bio-mathematical models of CRT in NSCLC (21) and evolutionary TKI resistance in advanced EGFR mutant NSCLC (11). An exponential vector system (See Methods, **Eqs. 1-14**) tracked the number of TKI resistant, persistent, and sensitive clonogenic cells (clonogens) and separated tumor burden into local versus distant compartments to deconvolve in-field locoregional failures and out of field distant failures, rather than simply modeling overall survival as a function of total tumor burden. This was an important distinction as post-progression treatment in the era of targeted therapy can be highly patient specific with varying degrees of response (22). Comorbidity related deaths were implemented into our model with a Monte Carlo Russian Roulette formalism (See Methods, **Eqs. 15-21**) and estimated from an analysis of the Surveillance, Epidemiology, and End Results (SEER) Program for a regional lung cancer population (23, 24). While a recent SEER analysis demonstrated that lung cancer patients receiving TKIs were less likely to be smokers, TKI recipients did not have a statistically significant difference the Charleson comorbidity index (0 vs. 1+) (25). A pictorial summary conceptualizing the model and its main endpoints is shown in **Fig. 1**.

**Figure 1:**
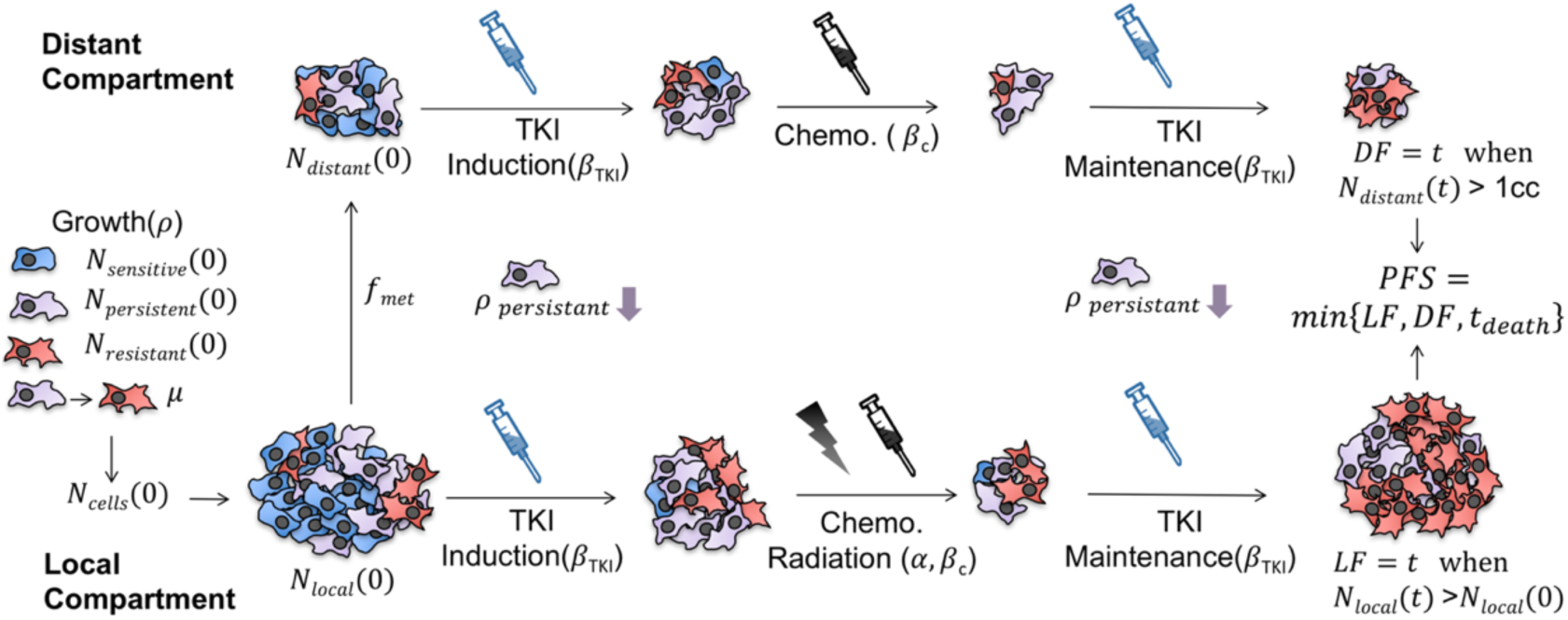
Graphical illustration of the distant and local tumor progression model. TKI sensitive, persistent, and resistant cells are shaded blue, purple, and red, respectively. The distant tumor compartment (top) is only susceptible to systemic agents, in this case chemotherapy and TKI therapy. The local tumor compartment (bottom) is additionally affected by radiation therapy. Note that TKI therapy is not only specific to sensitive cells but also leads to slowed growth of persistent cells. Chemo-radiation however is assumed to have equal effects on each cell subpopulation. The initial number of distant cells is assumed to be a constant fraction of the cells in the initial primary tumor.

### Model Calibration for Predicting Local and Distant Recurrence Dynamics of NSCLC

While the tumor progression model tracked a single tumor volume trajectory, a general patient population with heterogeneity in their presentation, response to treatment, and recurrence patterns was simulated by creating distributions for the model parameters. These distributions were then randomly sampled for each run (i.e. simulated patient) of the tumor progression model, yielding a histogram of times to events, which in turn was used to create simulated Kaplan-Meier (K-M) curves for the distributed population (see Methods). The model parameter distributions were then optimized such that the model predicted K-M curves matched clinically reported K-M curves, as was done in Geng et al. (21). In this work, the growth and radiosensitivity distributions were fitted to literature K-M curves of freedom from local and distant failure (FFLF and FFDF) in wildtype (WT) and EGFR+ locally advanced NSCLC populations receiving definitive concurrent CRT (22, 26, 27). The TKI model parameters were derived from a model-based analysis of advanced EGFR+ NSCLC patients (see Methods). A full summary of the optimized model parameters along with the source of the calibration data is shown in **Table 1**.

**Table 1.** Parameter Overview: Overview of parameters along with data sources used for model calibration and testing.

| Key Model Parameters used in Tumor Progression Model |  |  |  |
| --- | --- | --- | --- |
| Parameter | Description | Value | Source |
| $N_{cells}(0)$ | initial number of local clonogens | $[8.2 \cdot 10^6 - 6.7 \cdot 10^{11}]$ [cells] | Geng et al.<sup>21</sup> |
| $K$ | gompertzian carrying capacity | $8.2 \cdot 10^{12}$ [cells] | Geng et al.<sup>21</sup> |
| $V_p(0) = N_{persistent}(0)/N_{cells}(0)$<br>$V_r(0) = N_{resistant}(0)/N_{cells}(0)$ | initial fraction of TKI persistent & TKI resistant clonogens | $[0.05 - 0.5]$ [-]<br>$[10^{-4} - 10^{-1}]$ [-] | Grassberger et al.<sup>11</sup> |
| $\mu$ | resistant mutation probability | $10^{-7}$ [-] | Grassberger et al.<sup>11</sup> |
| $\beta_{TKI}$ | TKI cell kill parameter | log-norm ($\mu=2, \sigma=7$) $[\text{mg}/\text{m}^2]^{-1}$ ; ($>1$) | this work; Grassberger et al.<sup>11</sup> |
| $\beta_c$ | chemotherapy cell kill parameter | log-norm ($\mu=.028, \sigma=6.8 \cdot 10^{-4}$) $[\text{mg}/\text{m}^2]^{-1}$ | Geng et al.<sup>21</sup> |
| $f_{met}$ | initial number of metastatic clonogens as a fraction of $N_{cells}(0)$ | $2 \cdot 10^{-6}$ [-] | this work |
| $\rho$ | NSCLC growth rate | log-norm ($\mu=6 \cdot 10^{-5}, \sigma=.0055$) $[\text{day}]^{-1}$ | this work; Geng et al.<sup>21</sup> |
| $\alpha$ | WT NSCLC radiosensitivity | log-norm ($\mu=.10, \sigma=.17$) $[\text{Gy}]^{-1}$ | this work; Geng et al.<sup>21</sup> |
| $\alpha_{EGFR}$ | EGFR+ NSCLC radiosensitivity | log-norm ($\mu=.16, \sigma=.32$) $[\text{Gy}]^{-1}$ | this work; Geng et al.<sup>21</sup> |
| $t_{1/2}$ | chemotherapy plasma half life | 24 [hr] | Geng et al.<sup>21</sup> |
| $\kappa$ | TKI plasma decay factor | $0.0465$ $[\text{hr}]^{-1}$ | Grassberger et al.<sup>11</sup>, Foo et al.<sup>65</sup> |
| $r_{death}$ | comorbidity death rate for regional lung cancer | $0.55$ [%/mo] | this work; SEER<sup>23-24</sup> |
| Clinical Data used for Calibrating & Testing Model |  |  |  |
| Dataset | Description | Endpoint | Source |
| Calibration (N=118, USA) | institutional LF & DF rates in EGFR+/WT LA-NSCLC | FFLF & FFDF [mo] | Mak et al.<sup>27</sup> |
| Calibration (N=95, Japan) | institutional LF & DF rates in EGFR+/WT LA-NSCLC | FFLF & FFDF [mo] | Lim et al.<sup>26</sup> |
| Calibration (N=185, S. Korea) | institutional LF & DF rates in EGFR+/WT LA-NSCLC | FFLF & FFDF [mo] | Yagishita et al.<sup>22</sup> |
| Testing (N=86, France, Italy, Spain) | phase III clinical trial of TKIs in metastatic NSCLC (NCT00446225) | PFS [mo] | EURTRAC Trial, Rosell et al.<sup>1</sup> |
| Testing (N=598, Worldwide) | phase III clinical trial of concurrent chemoradiation in locally advanced NSCLC (NCT00686959) | PFS [mo] | PROCLAIM Trial, Senan et al.<sup>29</sup> |
| Testing (N=6 Clinical Trials) | meta-analysis of 6 clinical trials of sequential versus concurrent chemoradiotherapy in LA-NSCLC | FFLF & FFDF [mo] | Auperin et al.<sup>30</sup> |

While the 2 yr. FFLF for wild-type (WT) NSCLC has been shown to be relatively poor in the range of [54-63] %, EGFR-mutant tumors have a noticeably better response with a 2 yr. FFLF in the range [70-87] % (22, 26, 27). This differential response in FFLF was observed after CRT alone and the results were not confounded by TKI administration. Furthermore, *in-vitro* studies have demonstrated an enhance radiosensitivity of EGFR mutant NSCLC compared to WT (28), and so in our model unique radiosensitivity distributions were defined for WT and EGFR mutant populations separately and optimized against FFLF. Distant metastases have been observed to be the most common form of recurrence with a 2 yr. FFDF in the range of [31-43] % with no statistically significant relationship to EGFR status (22, 26, 27). Therefore, a common metastatic fraction and growth rate distribution were defined for both populations and optimized against FFDF, as shown in **Fig. 2**. For each calibration step, two parameters were optimized simultaneously, and while each two parameters exhibited a correlative relationship, a global solution was determined (**Suppl. Fig. 1**).

**Figure 2.**
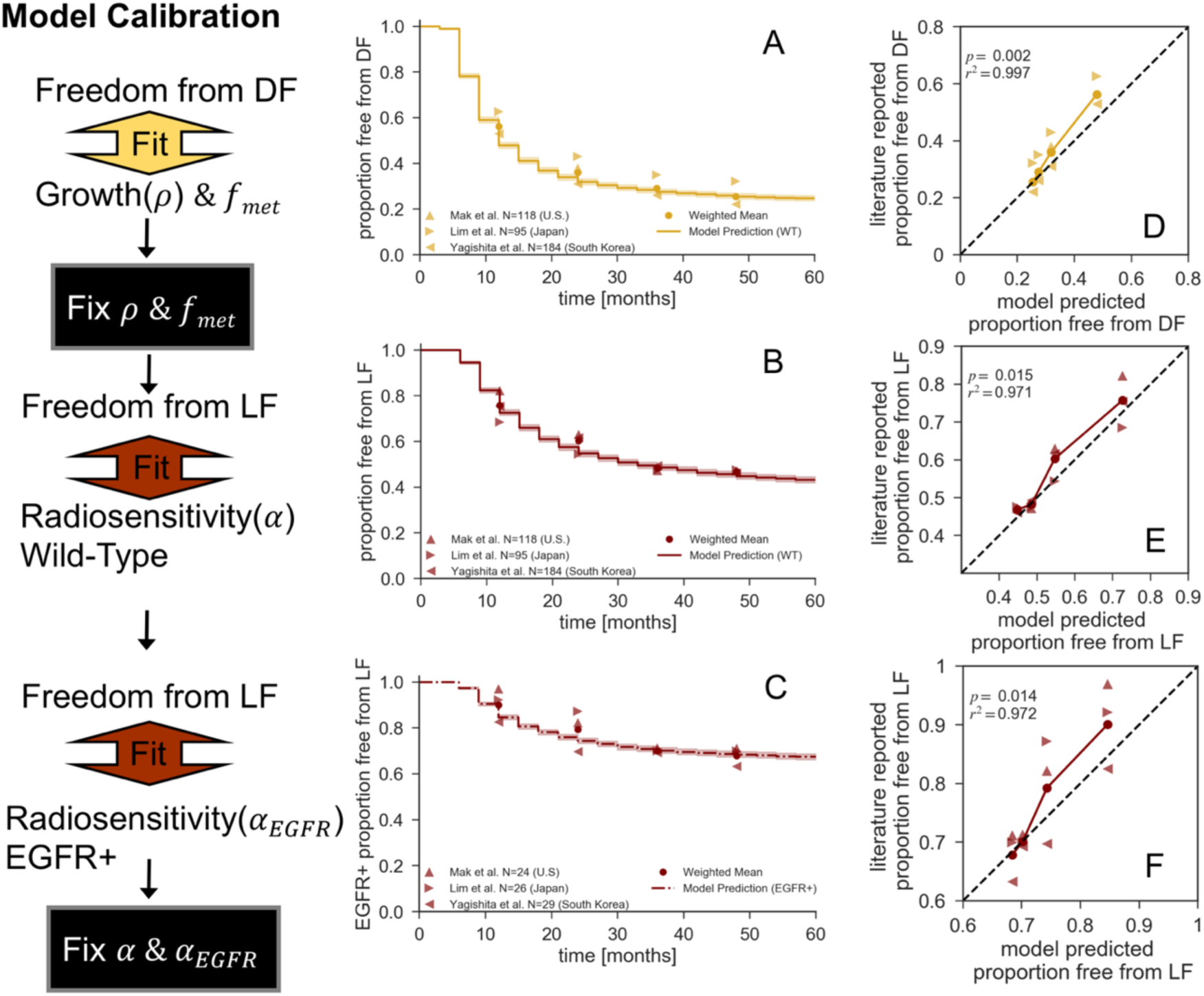
Model Calibration: A-C Simulated freedom from distant (A) and local (B, WT and C, EGFR+) failure Kaplan-Meier curves using the calibrated model parameters. D-F Corresponding model predicted versus literature reported failure rates at 1, 2, 3, and 4 yr. time points. The solid line represents the model predicted versus the weighted mean of literature reported failures, with corresponding linear regression and summary statistics. The black dashed line represents unity.

The model predicted FFDF curve for the optimal growth and metastatic fraction parameters is shown in **Fig. 2(A)** and exhibited a strong correlation to the weighted mean of the literature reported values (r^2^>0.99, p=0.002, **Fig. 2(D)**). Similarly, the model predicted FFLF curves with the optimized WT and EGFR+ radiation sensitivity distributions matched the literature reported values as shown in Fig. 2(B) and (C), respectively. For both populations, the model predicted FFLF were similarly strongly correlated to reported values (r^2^>0.97, p<0.02, Fig. 2(E) and (F)). Additionally, the model predicted PFS was calculated with the optimized parameters for WT and EGFR mutant populations, which yielded only a minor increase in PFS for the more radiosensitive EGFR mutant population (HR = 1.06 CI, 1.02-1.09, **Suppl. Fig. 2**). This was in accordance with literature reported observations of a lack of statistical significant difference in time to first recurrence between the WT and EGFR mutant populations (22, 27).

### Retrospective Model Validation Against Clinical Trial Outcomes

With all of the model parameters calibrated and fixed, the tumor progression model was then validated against independent data sets from multi-institutional phase 3 trials of either TKI alone or CRT alone (see Methods). First, the model predicted PFS in an advanced stage IV population was compared against the TKI arm of the EURTAC trial, which compared chemotherapy versus TKI in advanced EGFR mutant NSCLC (**Fig. 3(A)**) (1). A recent meta-analysis of first generation TKIs in EGFR+ advanced NSCLC found the EURTAC trial to have the most similar effect to the median of the six analyzed trials (HR EURTAC = 0.42, CI, 0.27-0.64, HR median = 0.37, CI, 0.27-0.52) making it an appropriate benchmarking dataset (17). The model predicted PFS was similar to the trial reported PFS (r^2^=0.92, p<1e-5, **Fig. 3(D)**), validating the rate of progression during TKI administration as predicted by the model. A waterfall plot of the maximum change in tumor volume from baseline is shown for a simulated patient population (n=256) in **Fig. 3(C)**, with each bar color-coded by degree of initial TKI resistance. From an analogous waterfall plot reported in the EURTAC trial, we see that the trial and the model simulated populations have similar tumor response dynamics with a median tumor volume decrease of 50%-75% and <10% of patients exhibiting progressive disease, demonstrating the validity of the derived TKI model parameters. Furthermore, the capacity of the model to stochastically embody a heterogeneous population is shown in **Fig. 3(C)**, as tumor shrinkage was not simply a function of pre-existing resistance but was modulated by other factors such as growth rate.

**Figure 3.**
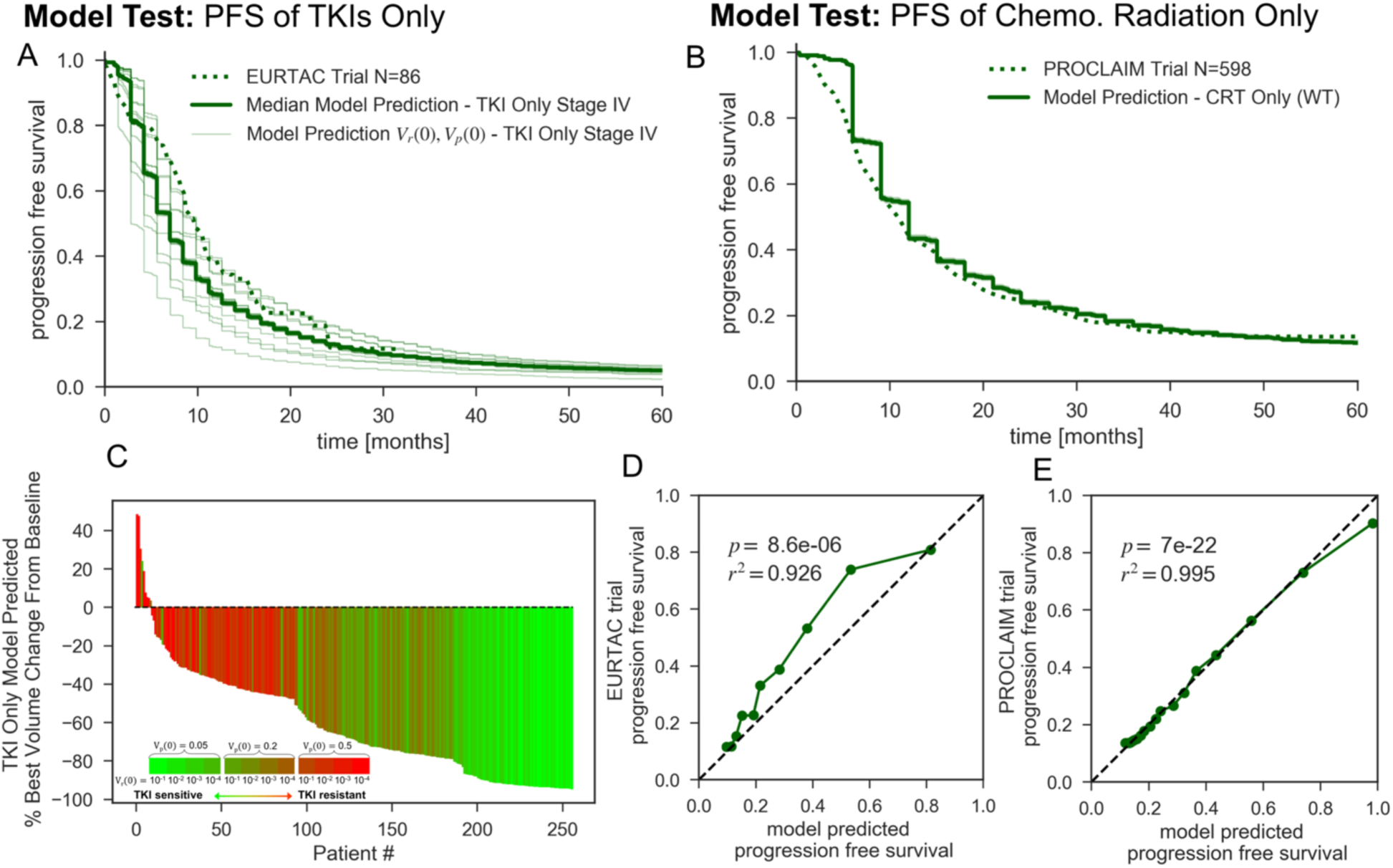
Model Validation: (A) Model predicted and trial reported PFS K-M curves for stage IV, EGFR+ NSCLC populations receiving TKIs until progression. The 12 thin K-M curves represent each initial starting persistent and resistant fraction combination (see Methods), while the full line is the median response. (B) Model predicted and trial reported PFS K-M curves for stage III NSCLC populations receiving definitive CRT. (C) Waterfall plot of best model predicted volume change from baseline for stage IV, EGFR+ NSCLC populations receiving TKIs until progression for 256 modeled patients, color-coded by the 12 initial TKI persistence/resistance conditions. Model predicted versus trial reported PFS over 3 month intervals are shown in (D) and (E) corresponding the K-M curves of (A) and (B), respectively. Linear regression summary statistics of model predicted versus trial reported PFS are displayed with the black dashed line representing unity.

Next, model predicted PFS for a WT LA-NSCLC population receiving concurrent CRT was compared against the results of the PROCLAIM trial (29), representing the pre-PACIFIC standard of care for CRT alone (**Fig. 3(B)**). Model predicted PFS for concurrent CRT correlated very strongly to the aggregated reported PFS of the trial (r^2^>0.99, p<1e-21, **Fig. 3(E)**), validating the absolute rate of recurrences during concurrent CRT as predicted by the model. Additionally, the model predicted local and distant failure dynamics for sequential versus concurrent CRT was in accordance with the results of a meta-analysis of six randomized trials comparing chemotherapy scheduling in LA-NSCLC (30). The model accurately predicted a significant benefit in local failure rates for concurrent CRT with a modeled HR of 0.79 (CI, 0.77-0.82), while the meta-analysis reported a HR of 0.77 (CI, 0.62-0.95, p=0.01) (**Suppl. Fig. 4(A)**). Furthermore, the model predicted no difference in distant failure rates for sequential versus concurrent CRT (HR = 1.00, CI, 0.97-1.03), which was consistent with the findings of the meta-analysis that there was no statistically significant effect of chemotherapy scheduling on distant failure rates (HR = 1.04, CI, 0.86-1.25, p=0.69) (**Suppl. Fig. 4(B)**). Together, these results provide strong evidence for the validity of the model predicted local and distant recurrence patterns during various treatment schedules as compared to current NSCLC multicenter clinical trials.

### Estimation of Improved Outcomes for TKI Induction and Maintenance

With the same simulated patient population and treatment parameters used during model validation (histograms shown in **Suppl. Fig. 3**), the expected recurrence dynamics of combining CRT and TKI therapy were explored. Two main treatment designs were simulated for locally advanced EGFR mutant NSCLC: TKI induction with daily administration up to 16 weeks followed by definitive concurrent CRT with and without adjuvant TKI maintenance. These two treatment schemes were chosen to approximate the format of ongoing combined TKI+CRT trial protocols (see Methods). TKI therapy concurrent with CRT was not modeled as initial clinical experience suggests the potential for increased toxicity and also a non-synergistic efficacy (31-33). The hypothesis explaining these results was that TKIs cause G1 cell-cycle arrest stunting cell replication antagonizing both chemotherapy and radiotherapy (34, 35), despite both agents having cytotoxic effects regardless of cell-division.

The predicted 2 yr., 3 yr., and 5 yr. FFLF, FFDF, and PFS as a function the TKI induction length with and without adjuvant TKI maintenance are plotted in **Suppl. Fig. 5**, while the endpoints over the entire design space are tabulated in **Suppl. Fig. 6**. For TKI induction without maintenance, the greatest predicted benefit for FFLF, FFDF, and PFS over CRT alone occurred with 2 wks. of induction (Δ5 yr. FFLF 1.5%, Δ5 yr. FFDF 11.1%, Δ5yr. PFS 6.5%), with a decreasing benefit as the induction length increased resulting in a similar predicted outcome to CRT alone with 16 wks. of induction. For each endpoint, there was a predicted additive benefit for including adjuvant TKI maintenance to induction regardless of the time point or duration of the induction period (**Suppl. Fig. 5**). But, the greatest benefit of including TKI maintenance was seen at shortest induction periods (Δ5yr. FFLF 5.8%, Δ5yr. FFDF 23.7%, Δ5yr. PFS 15.8% with 0 wks. induction and Δ5yr. FFLF 3.5%, Δ5yr. FFDF 7.7%, Δ5yr. PFS 6.7% with 2 wks. induction compared to CRT alone), with outcomes monotonically worsening as a function of induction length for each endpoint. The local failure rate was the least sensitive to the length of induction, given the high radiosensitivity and local control rates of EGFR+ NSCLC. The most dramatic effects were seen in the distant failure rates, which have an enhanced sensitivity to the evolutionary dynamics during induction as the subsequent chemotherapy was the only modeled therapeutic able to target the occult TKI resistant subpopulation in the distant compartment.

### Longer Induction is Predicted to Induce TKI Resistance

The maximum benefit observed with 2 wks. of TKI induction when maintenance was not administered is due to the fact that this time point presents a balance in benefit for both responders and non-responders: if a patient does not respond at all there is not much additional growth that early, while the responders will have at least derived some benefit through volume shrinkage. Furthermore, the benefit of delaying progression by means of extending the induction length (≥ 4 wks.) was gradually outweighed by the proliferation of the resistant population and resulting tumor volume increase prior to CRT. But surprisingly, when adjuvant maintenance was administered, the shortest induction periods were predicted to have the lowest local and distant failure rates. This suggests that the relative benefits of shrinking the tumor volume before CRT with TKIs was outweighed by the cost of acquired TKI resistance caused by targeted evolution during induction, which dramatically reduced the efficacy of TKI maintenance. This tradeoff in competing advantages of tumor size and TKI sensitivity is exemplified in the Kaplan-Meier analysis and simulated tumor volume trajectories displayed in **Fig. 4**.

**Figure 4.**
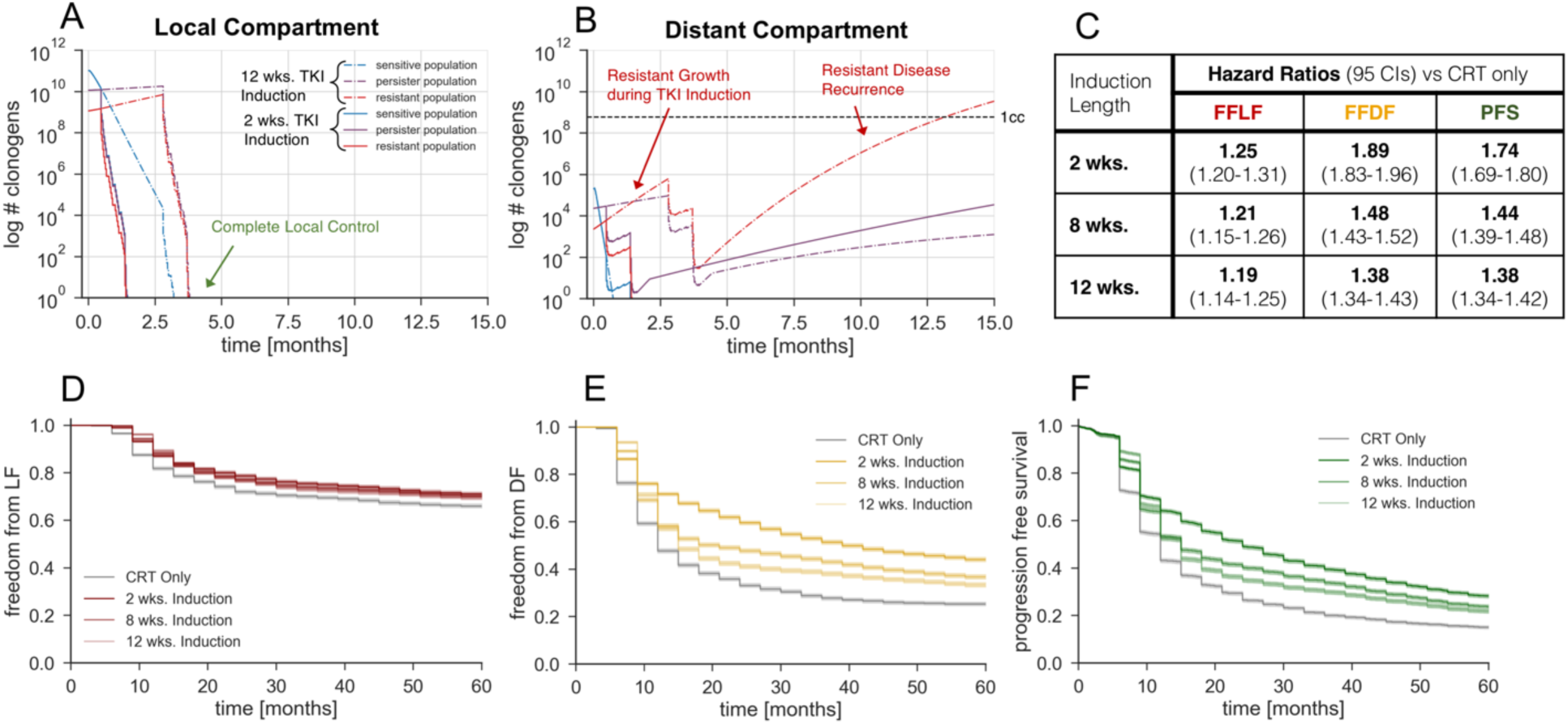
Model Predictions for Variable Induction Periods: Model predicted freedom from local failure (A), freedom from distant failure (B), and progression free survival (C) K-M curves for various induction lengths. The simulated treatment regimen is TKI induction, chemoradiotherapy, and adjuvant TKI maintenance. The local and distant tumor volume trajectory of the median simulated patient with an initial persistent fraction of 0.1 and initial resistant fraction of 0.01, stratified by TKI response cell subtypes, are shown in D and E respectively. Hazard ratios corresponding to the K-M curves (A, B, and C) are shown in F.

The median tumor volume trajectory stratified by TKI sensitive, persistent, and resistant cell subtypes receiving 2 and 12 wks. of TKI induction with adjuvant maintenance are plotted for the local compartment in **Fig. 4(A)** and the distant compartment in **Fig. 4(B)**. While both treatment schedules resulted in local control (**Fig. 4(A)**), a 2 wk. induction resulted in control of the initially microscopic resistant subpopulation of the distant compartment with chemotherapy during CRT and a slow regrowth of the persistent subpopulation during maintenance (**Fig. 4(B)**). But over the course of the longer 12 wks. of induction, the resistant subpopulation of the distant compartment out competed and outgrew the slowly proliferating persistent subpopulation, resulting in a rapid, resistant distant recurrence after ∼1 yr. of adjuvant TKI therapy (**Fig. 4(B)**), controlling for the same efficacy of chemotherapy during CRT. Additionally, the resistant growth during induction was accelerated in the distant compartment compared to the local compartment because of the resource advantage at lower cell numbers inherently modeled with Gompertzian growth (11).

Kaplan-Meier curves for the simulated FFLF, FFDF, and PFS for various TKI induction and maintenance schedules show the long term predicted benefit over CRT alone in Fig. 4(D), (E), and (F), respectively. While TKI induction and maintenance was predicted to have modest FFLF benefit over CRT alone, there was little stratification between different induction lengths (**Fig. 4(D)**). But for FFDF (**Fig. 4(E)**) and PFS (**Fig. 4(F)**), there was a much stronger effect with greater stratification between induction lengths. However, stratification between induction lengths was not observed until ∼1 yr. after the start of treatment, due to the time needed for distant cells to proliferate to an observable threshold after CRT. In this analysis, a schedule of no induction with CRT and adjuvant TKI maintenance was not considered despite having the best predicted outcomes, as some amount of induction with a measurable response in tumor volume is needed to warrant daily TKI administration after CRT until progression.

Calculated hazard ratios for 2 wks., 8 wks., and 12 wks. of TKI induction with adjuvant maintenance quantified the relative effect size of each multimodal treatment schedule compared to CRT alone, independent of the simulated sample size. As shown in **Fig. 4(C)**, FFDF had the most dramatic effect size, having a hazard ratio of 1.89 (CI, 1.83-1.96) for 2 wks. induction dropping to 1.38 (CI, 1.34-1.43) for 12 wks. induction. The expected statistical significance of these proposed treatment schema at clinically relevant sample sizes were stochastically investigated, as described in the next section.

### Statistical Significance and Power in a Model-Based Trial Design

Evaluating the statistical significance between the predicted failure rates of different treatment arms was a non-trivial task as an arbitrarily large number of patients can be simulated, which could result in statistical significance even when hazard ratios were very close to one. Therefore, we determined the probability of reaching a certain level of statistical significance as a function of the number of trial patients. We utilized the stochastic nature of the model by randomly sampling multiple iterations of simulated patients in order to yield an estimate of the false negative rate at lower sample sizes and the corresponding statistical power (see Methods Section). Doing so, our model based analysis not only yielded an expected magnitude of effect between two multimodal treatments but also quantified the probability of observing the effect in a population of a given size.

A simulated two armed clinical trial of 2 wks. versus 12 wks. of TKI induction with CRT and adjuvant TKI maintenance was modeled with FFDF as the endpoint, which has both the strongest effect (**Fig. 4(F)**) and is particularly relevant for future targeted therapy trials given the relatively high rates of local control in NSCLC with CRT for EGFR-mutant patients. A depiction of the simulated evolution of TKI resistance illustrating the hypothesized mechanism of benefit to a shorter induction period is shown in **Fig. 5(A)**. Modeled FFDF K-M curves displayed a constant magnitude of effect but wider confidence intervals with decreased sample size (**Fig. 5(B)**). A heatmap of the log-rank p-values testing the significance between the two arms over 1000 iterations revealed the variability in detecting the effect with fewer patients in **Fig. 5(C)**. This was further quantified in **Fig. 5(D)**, where the distributions of median FFDF in each arm have a consistent mean but increasing variance with decreasing sample size. Note that the simulation ran for 5 yrs. and so the peak at 60 months can be attributed to iterations where the median FFDF was not yet reached. The distribution of log-rank p-values between the arms for the 1000 iterations is shown in **Fig. 5(E)** along with the median p-values. The fraction of iterations reaching statistical significance of 0.05 was estimated to be the statistical power, and is plotted as function of sample size in **Fig. 5(F)**. Thus, this analysis has projected that a clinical trial would need 256 patients per arm for a power of 79%.

**Figure 5.**
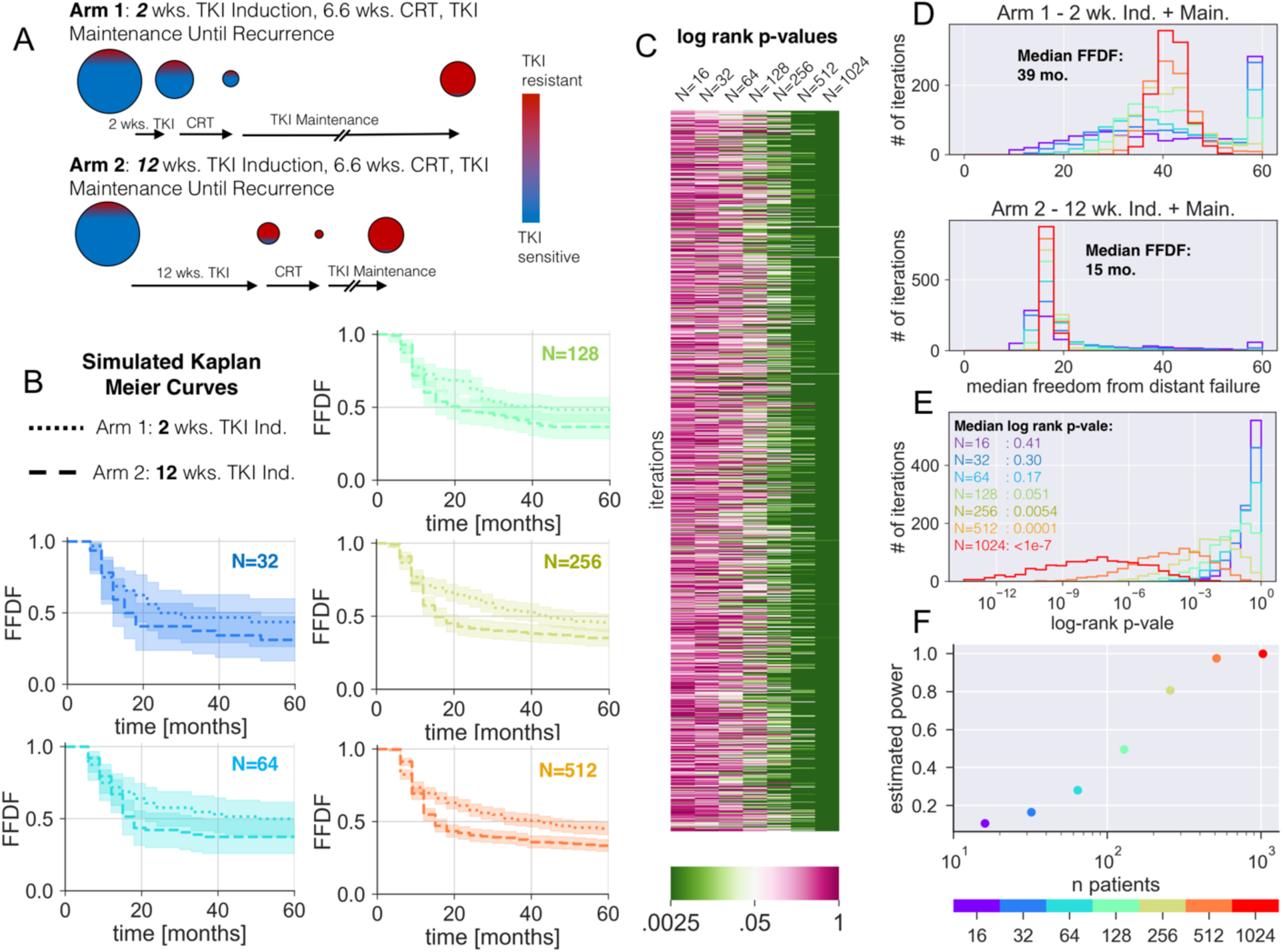
Simulated *in-silico* Induction Trial: (A) Illustration of the differential evolution of the TKI-resistant and sensitive populations for a given tumor burden between a short 2 wk. and long 12 wk. induction length. When CRT is done after a significant TKI induction period, the tumor shrinks with the targeted drug killing the TKI sensitive cells (blue), but with more TKI resistant (red) cells at the time of CRT, increasing the chance of a late TKI resistant recurrence if CRT isn’t curative. (B) Simulated FFDF K-M curves for 2 wk. versus 12 wk. induction lengths with increasing number of simulated patients. Each curve corresponds to the iteration with the median log-rank p-vale. (C) A heatmap of log rank p-values testing statistical difference between the 2 wk. versus 12 wk. FFDF K-M curves for 1000 iterations of the simulation at each sample size. (D) Histograms of the median FFDF for the 1000 iterations of the 2 wk. and 12 wk. induction simulations at each sample size. (E) Histogram of the log rank p-value between the 2 wk. versus 12 wk. induction simulations at each sample size. (F) Estimated statistical power as a function of sample size. Here statistical power was estimated as the fraction of iteration resulting in a p-value<0.05.

A corresponding analysis using PFS as the endpoint is displayed in **Suppl. Fig. 7**, which estimated that 512 patients per arm are needed to reach a power over 80%. While these analyses assumed a general population, the modeled population can be readily stratified into clinically distinguishable groups. For instance, modeled populations above and below the median initial tumor size (7.2 cm) will exhibit diverging distributions of predicted failure rates, which will consequently decrease and increase the required number of patients in each sub-population needed for an adequately powered clinical trial.

## Discussion

We have developed a mathematical framework to forecast the relative effectiveness of user-defined regimens of combined targeted-chemoradiation therapy and have applied this methodology to optimize the multimodal administration of tyrosine kinase inhibitors in locally advanced NSCLC. This framework integrates fundamental principles of tumor growth, radiation biology, and acquired drug resistance, with model parameters calibrated to institutional outcomes and predicted recurrence rates independently validated against clinical trial outcomes. We predict local and distant recurrence rates for various lengths of TKI induction before CRT and estimate the expected number of patients needed to observe a clinical benefit. In contrast to the current design of multimodal clinical trials which prescribe 2-3 month long induction periods, we discover an inverse relationship between progression and length of induction. In this sense no induction period at all would be optimal. However, some TKI induction is indicated because we need to see that the patient responds at all before proposing an adjuvant drug regimen, where response assessment is complicated by the preceding CRT regimen. We propose 2 weeks because we think one can observe a volumetric reduction at this point as measured by consistent volumetric segmentation.

After 2 weeks of TKI exposure, there may not be maximal macroscopic gross tumor volume change; however, at these early time points, the occult TKI resistant and persistent populations will still be microscopic and with greater potential for control by cytotoxic therapy. Even though we make the assumption that pre-existing resistant cells exist at therapy initiation, our result does not entirely depend on this assumption. Even assuming no pre-existing cells would favor shorter induction periods, as this reduces the probability of cells acquiring mutations that confer resistance. The sooner CRT is administered, the sooner the pool of possible persister cells that could acquire resistance is diminished, increasing the efficacy of the TKI maintenance regimen. This should also hold true for second and third generation TKIs as these drugs still result in acquired drug resistance through a combination of pre-existing and acquired mutations (5, 6), despite overcoming the EGFR T790M mutation which has indeed resulted in superior PFS compared to first generation TKIs (2)

Our result that shorter induction times lead to better outcomes contradicts expectations in current multimodal trial protocols for LA-NSCLC, but reflects the increasing risk of TKI resistance as a function of TKI exposure. Induction before CRT has several benefits, most notably decreasing tumor size, which has been shown to improve overall survival when treated with CRT (36, 37) and may also lead to surgical candidacy as outlined in the ASCENT trial protocol. But, upfront TKI exposure in advanced staged patients has resulted in a 50-75% average volume change (1), which is similar to the expected surviving fraction after a single 2 Gy radiation fraction (SF2Gy) in NSCLC (38). Additionally, upfront TKI exposure will extend the duration of treatment compared to CRT alone, which will ostensibly delay disease progression. But as patients systemically become resistant over the course of months, adjuvant TKI maintenance therapy may be rendered ineffective. Conversely, with upfront chemoradiation therapy and the ability to control TKI resistant cells, the adjuvant TKI maintenance has the potential to dramatically slow tumor regrowth. And in fact, a retrospective analysis of the treatment of brain metastases originating from EGFR mutant NSCLC demonstrated that upfront radiotherapy with adjuvant TKIs nearly doubled overall survival compared to upfront TKIs until progression and subsequent radiotherapy (34.1 vs. 19.4 mo.; p = .01) (39). But in the locally advanced setting, some initial TKI exposure is needed in order to demonstrate tumor response before maintenance therapy can be considered. Furthermore, there is evidence to suggest that TKI exposure may prime the cells for radiotherapy by inducing senescence(40)

Genetic heterogeneity present throughout a patients’ tumor predicates the clinically observable acquired treatment resistance (41). There are two main models explaining the mechanism by which intratumoral heterogeneity and consequently treatment resistance arises: (1) branched clonogenic evolution where subpopulations arise from an accumulation of mutations and (2) cancer stem cells (CSCs) that are predominantly dormant and treatment resistant, but uniquely give rise to the diversity of differentiated cancer cells (7, 42). Additionally, models have been proposed unifying CSCs and clonogenic evolution, suggesting that CSCs themselves can acquire favorable mutations leading to competing subpopulations (43). In this work, a form of clonogenic evolution was assumed with CSCs not explicitly modeled, as the focus of this analysis was acquired TKI resistance, which has been shown to occur through specific genetic mutations (44-46). As such, in our model TKI resistant cells are assumed to have the same chemo- and radiosensitivity as TKI sensitive cells, under the assumption that both subtypes have a similar phenotype to EGFR mutant NSCLC, which is notably sensitive to conventional CRT, which is in line with experimental evidence (28).

While our model does not explicitly account for the constricting effect of TKIs on vasculature, resource deprivation is implicitly modeled through the slowed growth of the TKI persistent subpopulations resulting in apparent stunted tumor growth. Future work could incorporate the effect of vasculature modulation into the model, by both EGFR-TKIs and also anti-angiogenic vascular endothelial growth factor inhibitors (VEGF-TKIs) which have shown great synergy when administered in combination (47). Chemotherapy or radiotherapy concurrent with TKIs was not considered in this study, but intercalated multimodal therapy has been shown to be feasible and the effects of which could be investigated in future studies (48). Furthermore, while we only considered the chemotherapy regimen used in RTOG 9410, clinical experience has demonstrated similar outcomes with a variety of chemotherapy regimens (49).

Based on our framework, future modeling studies might aim at stratifying simulated populations (e.g. high or low radiosensitivity *α*, tumor growth *ρ*) and determine which patient populations experience the most benefit from shorter versus longer induction periods, thus further personalized therapy. Potential biomarkers for NSCLC radiosensitivity demonstrated to be associated with survival outcomes are rad51 expression (50) and the polygenic radiosensitivity index (RSI) (51, 52). Additionally, ki67 expression and fluorothymidine uptake imaged by positron emission tomography have both shown to be related to tumor cell proliferation and could be potential biomarkers of tumor growth rate (53-56).

In conclusion, we have provided a mathematical framework for optimizing multimodal targeted therapy and provide an evolutionary argument against longer induction periods, as they could trigger the process of acquired drug resistance and may limit the efficacy of adjuvant therapy, possibly resulting in worse outcomes. According to our model a shorter induction period of 1-2 weeks had a greater chance of controlling TKI resistant cells with CRT, resulting in longer predicted progression free survival. Finally, the probability of observing a statistically significant increase in PFS due to a shorter induction period was stochastically derived as function of trial size, using randomly sampled heterogeneous patient populations. These model predictions are hypothesis generating and could have impact on clinical trial design. While this study has focused on optimizing TKI administration in combination with CRT for EGFR mutant NSCLC, the generalized framework outlined in this paper can be applied to oncogene-driven multimodal therapy designs in other cancers.

## Methods

### Mathematical Implementation of Local versus Distant Tumor Progression Model

We described the evolution of the number of TKI resistant (*N*_*R*_), persistent (*N*_*P*_), or sensitive (*N*_*S*_) clonogens in the distant (*N*_*D*_) or local (*N*_*L*_) compartments (**Eqs. 5-10**) with exponential factors accounting for the differential terms of Gompertzian cell growth, Norton-Simon cell kill by TKIs, log cell kill by chemotherapy, and linear-quadratic (LQ) radiation cell kill (with the assumption of *α*/*β* = 10). While older reports suggested *α*/*β* could be higher than 10 for lung cancer (57), our results were insensitive to the exact value of *α*/*β* as different fractionation schema were not considered. The initial local cell number was based on tumor volume distributions for each stage of NSCLC and an assumed cell density of 5.8×10^8^ cells/cm^3^ estimated from previous model based work of NSCLC tumor growth (21, 58, 59). The initial distant cell number was assumed to be a scalar fraction of the initial local cell number (**Eq. 2**), with distant failure defined as previously occult metastasis reaching a volume 1 cm^3^ (60) and a local failure defined as growth past the size at the start of treatment. Throughout treatment and regrowth, there was also a transitional term to account for acquired mutations from TKI persistent to resistant cells with the mutation rate *μ* assumed to be 10^−7^ based upon in-vitro studies of TKI resistance (61). The model also assumed that the growth of the persistent compartment was slowed during TKI administration as seen clinically (**Eq. 13**) (11, 44). The first order pharmacokinetic model of chemotherapy (*C*_*C*_, **Eq. 12**) and TKI plasma concentrations (*C*_*TKI*_, **Eq. 14**) used in our model were fully described in previous publications (11, 21). The state equations were implemented with 0.1 day resolution (Δ*t*) with the differential effect of each term calculated independently. Discrete treatment events, such as radiation delivery, were assumed to occur instantaneously at a single timepoint (**Eq. 11**). At a discrete time step, if a cell compartment went below 1 cell, it was assumed to be controlled and set to 0.

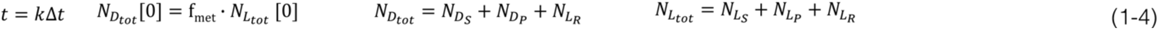

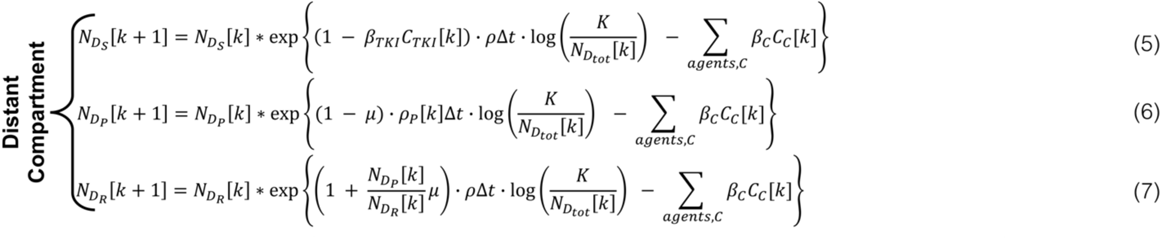

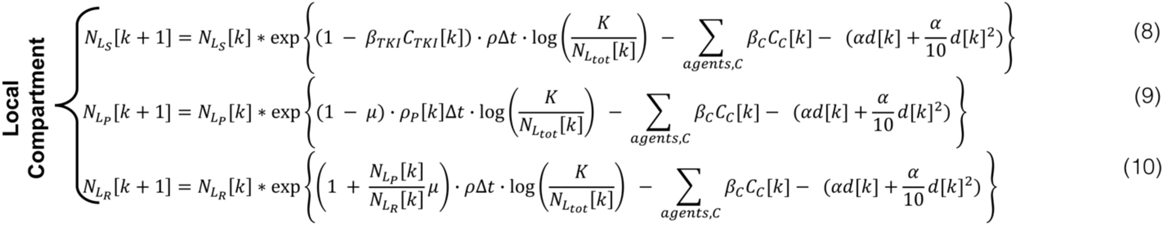

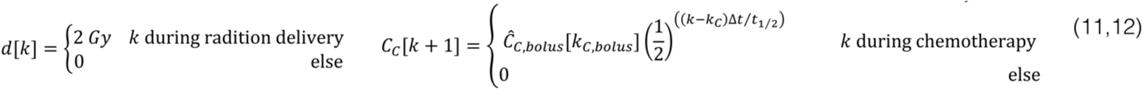

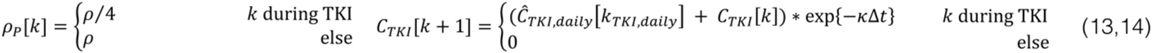

### Implementation of Deaths from Comorbidities and Failure Rates

To accurately model progression free survival (PFS) in a lung cancer population for which comorbidities can be significant, a non-cancerous death rate from comorbidities must be considered. Non-cancer related death rates during the 5 yrs. after CRT for lung cancer patients were retrieved from an analysis of the Surveillance, Epidemiology, and End Results (SEER).

This analysis stratified the 5 yr. survival probability, death probability from locally-advanced lung cancer, and non-cancerous death probability by age, sex, and severity of comorbidity status. The overall 5 yr. survival probability, cancerous death probability, and non-cancer death probability for the population was calculated to be 15.2%, 71.3%, and 13.5% respectively, given the patient age and sex frequency in the SEER analysis (23) and also the frequency of each comorbidity severity level reported in a separate analysis of SEER data (24). With the current standard of care including PET staging, concurrent chemotherapy, and intensity modulated radiotherapy, the current 5 yr. survival probability from locally-advanced lung cancer was approximated to be 28.8% (averaged 5 yr. overall survival from each arm of the recent PROCLAIM concurrent CRT trial). Assuming the same proportion of deaths from lung cancer and non-cancerous comorbidities but with this same survival probability, the adjusted 5 yr. death probability from cancer and comorbidities was estimated as 59.9% and 11.3%, respectively. The monthly death rate from comorbidities, *r*_*death*_, was then estimated to be 0.55%. The probability of dying from comorbidities on a given discrete month, *t*_*month*_ ∈ ℕ, and the cumulative probability of dying at *t*_*death*_ on or before *t*_*month*_, were be then calculated by monthly compounding of *r*_*death*_ (**Eqs. 15-16).** This cumulative distribution function was then inverted and sampled by a drawing a random number, *ζ* ∈ [0,1], to generate a single, stochastic drawing of the time of death by comorbidity *t*_*death*_ ∈ [0, ∞] for each simulated patient **(Eqs. 17-18**). Finally, the time to progression for an individual patient (**Eq. 21**) was defined as the earliest of time of local failure (*LF*, **Eq. 19**), time of distant failure (*DF* **Eq. 20**), or time of death from comorbidity (*t*_*death*_, **Eq. 18**). Note that *LF* and *DF* were assumed to be a multiple of 3 months since the start of treatment, in order to mimic the clinical practice post-treatment evaluation with CT approximately every 3 months.

### Hazard Ratio Calculation & Kaplan Meier Analysis

The *lifelines v0.18* python package was used for creating K-M curves and corresponding 95% confidence intervals. The input to the K-M estimator was a list of time to events (local failure, distant failure or progression) of length N, where N is the number of simulated patients. For the K-M analysis of the simulated data, patients were assumed to have all started treatment simultaneously with no censorship of patients. Hazard ratios between two population endpoints receiving different treatment schemes were calculated using the Mantel-Haenszel method (62), using the events table of each arm reported by the *lifelines v0.18* package.

### CRT Model Parameter Calibration

The log-normal model parameter distributions published by Geng et al. (21) were used, with the growth and radiosensitivity distributions recalibrated to three single institutional reports of freedom from local and distant failure (FFLF and FFDF) rates in wildtype (WT) and EGFR mutant locally advanced (LA) NSCLC populations receiving definitive concurrent CRT (22, 26, 27). In this analysis, WT specifically refers to a randomly sample from the general population not screened for EGFR status, with reported incidences of EGFR mutations of 24/118 (20%, US) (27), 26/95 (27%, Japan) (22), 29/184 (16%, South Korea) (26) and a mean of 79/397 (20%). The initial stage distribution of the simulated LA-NSCLC population was assumed to be 47% for Stage IIIA and 53% for IIIB as reported in a Phase 3 CRT trial (PROCLAIM) (29). Accordingly, the modeled concurrent CRT treatment was 2 Gy per fraction in 33 fractions of RT (per PROCLAIM) (29) with concurrent chemotherapy (per RTOG 9410 Arm 2) (63) as in our previous work. Optimization was performed where the normalized squared summed difference between the model predicted and the mean literature obtained FFDF or FFLF at 1, 2, 3, and 4 yrs. was calculated over the entire clinically relevant parameter space. For each calibration, five runs with n=1024 simulated patients were performed and aggregated at each parameter value to determine the global optimum (**Suppl. Fig. 1**). The standard deviation of the growth distribution, *ρ*_*σ*_, and the metastatic fraction, f_*met*_, were simultaneously optimized over the FFDF. These parameters were then fixed, and the mean and standard deviation of WT and EGFR mutant radiosensitivity distributions, *α*_*μ*_ and *α*_*σ*_, were simultaneously optimized for the WT and EGFR mutant FFLF, respectively.

For both the model calibration and validation, data was extracted from the literature reported K-M curves with the *WebPlotDIgitzer V4.2* program (64). Additionally, the relationship between the model predicted versus literature reported failure rates were quantified with a linear regression, implemented with *linregress* function in the *SciPy* library.

### TKI Model Parameter Calibration

The TKI model parameters for our simulated population were derived from a previous model based analysis of n=20 tumor volume trajectories in advanced EGFR mutant NSCLC patients receiving erolitnib (11). Initial persistent fractions of [0.05, 0.2, 0.5] and an initial resistance fractions of [10^−4^, 10^−3^, 10^−2^, 10^−1^] were modeled in equal proportion, representing a range of initial TKI sensitivities observed in the patient specific tumor volume trajectories and in an in-vitro clonogenic assay of cells derived from human NSCLC solid tumors (44). Similarly, the TKI cell kill distribution was estimated to be log-normal with a mean of 2 and a standard deviation of 7, such that the median and maximum TKI cell kill value approximately matched those estimated from the patient specific tumor volume trajectories (patient specific median = 7.8 versus median of log-normal = *e*^*μ*^= *e*^2^ = 7.4; patient specific maximum = 20 versus log-norm 99.9^th^ percentile = 21 for n = 1024 simulated patients).

### Model Validation Against Clinical Trials

For the model validation against phase 3 clinical trials of CRT alone and TKIs alone, n=1024 patients were simulated having initial persistent fractions of [0.05, 0.2, 0.5] and initial resistance fractions of [10^−4^, 10^−3^, 10^−2^, 10^−1^] for a total of n=1024×4×3=12288 patients. Histograms of the used model parameters are shown in **Suppl. Fig. 3**. The modeled concurrent CRT treatment was implemented the same as in the model calibration (per PROCLAIM and per RTOG 9410 Arm 2) while the sequential CRT was implemented per RTOG 9410 Arm 1 with the chemotherapy preceding the radiotherapy (63), resulting in the distant compartment receiving the same treatment in both schedules. Both concurrent chemotherapy arms of the PROCLAIM trial were aggregated as no effect was observed. For the model validation against TKIs alone, the TKI treatment was implemented similar to a previous report (11, 65), wherein daily bolus of erlotinib was administered within a first order pharmacokinetic model resulting in a daily plasma concentration of the drug (**Eq. 14)**. The daily administration continues until progression, as was done in the comparative trials. Additionally, the advanced (stage IV) tumor size distribution from Geng et al. (21) was assumed given the comparative trial was similarly performed in an advanced population. Finally, both the simulated tumor size change from baseline and recurrences were measured every six weeks in accordance with the TKI monotherapy trail.

### Multimodal Target-Chemoradiotherapy Treatment Modeling

The two main treatment designs simulated in locally advanced EGFR mutant NSCLC were TKI induction with daily administration up to 16 weeks, followed by definitive concurrent CRT, with and without adjuvant TKI maintenance until a distant or local tumor recurrence. The concurrent CRT was modeled the same as during the model calibration and validation. These two treatment schemes are very similar to ongoing combined TKI+CRT trial protocols (NCT 01822496, 12 wks. TKI induction followed by definitive CRT and NCT 01553942, 8 wks. TKI induction followed by definitive CRT, followed by surgery if possible, followed by 6 wks. chemotherapy, and finally adjuvant TKI maintenance if there was initial response), with the exception that our model does not take into account surgery.

### Statistical Power Estimation

To analyze the statistical significance of differences in the endpoints between various induction lengths, a Monte Carlo based power calculation was performed to simulate several iterations of a two-arm trial. A short (2 wk.) and a long (12 wk.) induction length arm were simulated with 7 wks. concurrent CRT and adjuvant TKI therapy with n_i_ = [16, 32, 64,128, 256, 512,1024] patients in each arm. This was iterated 1000 times by randomizing all N=12288 patients using the *numpy.random.shuffle* routine and again analyzing the first n_i_ patients. For each iteration of each sample size n_i_, the median PFS or FFDF of each arm was recorded. The difference in rate of progression or distant failure between the short and long induction lengths was determined to be statistically significant by a log-rank test, which was implemented using the *lifelines v0.18* python package. The power as function of sample size was then estimated as the fraction of the 1000 iterations which did reach statistical significance (p < 0.05).

## References

1. Rosell R, Carcereny E, Gervais R, Vergnenegre A, Massuti B, Felip E, Palmero R, Garcia-Gomez R, Pallares C, Sanchez JM, Porta R, Cobo M, Garrido P, Longo F, Moran T, Insa A, De Marinis F, Corre R, Bover I, Illiano A, Dansin E, de Castro J, Milella M, Reguart N, Altavilla G, Jimenez U, Provencio M, Moreno MA, Terrasa J, Munoz-Langa J, Valdivia J, Isla D, Domine M, Molinier O, Mazieres J, Baize N, Garcia-Campelo R, Robinet G, Rodriguez-Abreu D, Lopez-Vivanco G, Gebbia V, Ferrera-Delgado L, Bombaron P, Bernabe R, Bearz A, Artal A, Cortesi E, Rolfo C, Sanchez-Ronco M, Drozdowskyj A, Queralt C, de Aguirre I, Ramirez JL, Sanchez JJ, Molina MA, Taron M, Paz-Ares L, Spanish Lung Cancer Group in collaboration with Groupe Francais de P-C, Associazione Italiana Oncologia T. Erlotinib versus standard chemotherapy as first-line treatment for European patients with advanced EGFR mutation-positive non-small-cell lung cancer (EURTAC): a multicentre, open-label, randomised phase 3 trial. Lancet Oncol. 2012;13(3):239–46. doi: 10.1016/S1470-2045(11)70393-X. PubMed PMID: 22285168.

2. Soria JC, Ohe Y, Vansteenkiste J, Reungwetwattana T, Chewaskulyong B, Lee KH, Dechaphunkul A, Imamura F, Nogami N, Kurata T, Okamoto I, Zhou C, Cho BC, Cheng Y, Cho EK, Voon PJ, Planchard D, Su WC, Gray JE, Lee SM, Hodge R, Marotti M, Rukazenkov Y, Ramalingam SS, Investigators F. Osimertinib in Untreated EGFR-Mutated Advanced Non-Small-Cell Lung Cancer. N Engl J Med. 2018;378(2):113–25. doi: 10.1056/NEJMoa1713137. PubMed PMID: 29151359.

3. Holohan C, Van Schaeybroeck S, Longley DB, Johnston PG. Cancer drug resistance: an evolving paradigm. Nat Rev Cancer. 2013;13(10):714–26. doi: 10.1038/nrc3599. PubMed PMID: 24060863.

4. Turner NC, Reis-Filho JS. Genetic heterogeneity and cancer drug resistance. Lancet Oncol. 2012;13(4):e178–85. doi: 10.1016/S1470-2045(11)70335-7. PubMed PMID: 22469128.

5. Chabon JJ, Simmons AD, Lovejoy AF, Esfahani MS, Newman AM, Haringsma HJ, Kurtz DM, Stehr H, Scherer F, Karlovich CA, Harding TC, Durkin KA, Otterson GA, Purcell WT, Camidge DR, Goldman JW, Sequist LV, Piotrowska Z, Wakelee HA, Neal JW, Alizadeh AA, Diehn M. Circulating tumour DNA profiling reveals heterogeneity of EGFR inhibitor resistance mechanisms in lung cancer patients. Nat Commun. 2016;7:11815. doi: 10.1038/ncomms11815. PubMed PMID: 27283993; PMCID: PMC4906406.

6. Ortiz-Cuaran S, Scheffler M, Plenker D, Dahmen L, Scheel AH, Fernandez-Cuesta L, Meder L, Lovly CM, Persigehl T, Merkelbach-Bruse S, Bos M, Michels S, Fischer R, Albus K, Konig K, Schildhaus HU, Fassunke J, Ihle MA, Pasternack H, Heydt C, Becker C, Altmuller J, Ji H, Muller C, Florin A, Heuckmann JM, Nuernberg P, Ansen S, Heukamp LC, Berg J, Pao W, Peifer M, Buettner R, Wolf J, Thomas RK, Sos ML. Heterogeneous Mechanisms of Primary and Acquired Resistance to Third-Generation EGFR Inhibitors. Clin Cancer Res. 2016;22(19):4837–47. doi: 10.1158/1078-0432.CCR-15-1915. PubMed PMID: 27252416.

7. Altrock PM, Liu LL, Michor F. The mathematics of cancer: integrating quantitative models. Nat Rev Cancer. 2015;15(12):730–45. doi: 10.1038/nrc4029. PubMed PMID: 26597528.

8. Grassberger C, Scott JG, Paganetti H. Biomathematical Optimization of Radiation Therapy in the Era of Targeted Agents. Int J Radiat Oncol Biol Phys. 2017;97(1):13–7. doi: 10.1016/j.ijrobp.2016.09.008. PubMed PMID: 27979444.

9. Foo J, Michor F. Evolution of acquired resistance to anti-cancer therapy. J Theor Biol. 2014;355:10–20. doi: 10.1016/j.jtbi.2014.02.025. PubMed PMID: 24681298; PMCID: PMC4058397.

10. Bozic I, Reiter JG, Allen B, Antal T, Chatterjee K, Shah P, Moon YS, Yaqubie A, Kelly N, Le DT, Lipson EJ, Chapman PB, Diaz LA, Jr., Vogelstein B, Nowak MA. Evolutionary dynamics of cancer in response to targeted combination therapy. Elife. 2013;2:e00747. doi: 10.7554/eLife.00747. PubMed PMID: 23805382; PMCID: PMC3691570.

11. Grassberger C, McClatchy DM, Geng C, Kamran SC, Fintelmann F, Maruvka YE, Piotrowska Z, Willers H, Sequist LV, Hata AN, Paganetti H. Patient-specific tumor growth trajectories determine persistent and resistant cancer cell populations during treatment with targeted therapies. Cancer Research. 2019:canres.3652.2018. doi: 10.1158/0008-5472.CAN-18-3652.

12. Yoon N, Vander Velde R, Marusyk A, Scott JG. Optimal Therapy Scheduling Based on a Pair of Collaterally Sensitive Drugs. Bull Math Biol. 2018;80(7):1776–809. doi: 10.1007/s11538-018-0434-2. PubMed PMID: 29736596.

13. Nichol D, Rutter J, Bryant C, Hujer AM, Lek S, Adams MD, Jeavons P, Anderson ARA, Bonomo RA, Scott JG. Antibiotic collateral sensitivity is contingent on the repeatability of evolution. Nat Commun. 2019;10(1):334. doi: 10.1038/s41467-018-08098-6. PubMed PMID: 30659188; PMCID: PMC6338734.

14. Leder K, Pitter K, LaPlant Q, Hambardzumyan D, Ross BD, Chan TA, Holland EC, Michor F. Mathematical modeling of PDGF-driven glioblastoma reveals optimized radiation dosing schedules. Cell. 2014;156(3):603–16. doi: 10.1016/j.cell.2013.12.029. PubMed PMID: 24485463; PMCID: PMC3923371.

15. Chmielecki J, Foo J, Oxnard GR, Hutchinson K, Ohashi K, Somwar R, Wang L, Amato KR, Arcila M, Sos ML, Socci ND, Viale A, de Stanchina E, Ginsberg MS, Thomas RK, Kris MG, Inoue A, Ladanyi M, Miller VA, Michor F, Pao W. Optimization of dosing for EGFR-mutant non-small cell lung cancer with evolutionary cancer modeling. Sci Transl Med. 2011;3(90):90ra59. doi: 10.1126/scitranslmed.3002356. PubMed PMID: 21734175; PMCID: PMC3500629.

16. Yu HA, Sima C, Feldman D, Liu LL, Vaitheesvaran B, Cross J, Rudin CM, Kris MG, Pao W, Michor F, Riely GJ. Phase 1 study of twice weekly pulse dose and daily low-dose erlotinib as initial treatment for patients with EGFR-mutant lung cancers. Ann Oncol. 2017;28(2):278–84. doi: 10.1093/annonc/mdw556. PubMed PMID: 28073786; PMCID: PMC5834093.

17. Gao G, Ren S, Li A, Xu J, Xu Q, Su C, Guo J, Deng Q, Zhou C. Epidermal growth factor receptor-tyrosine kinase inhibitor therapy is effective as first-line treatment of advanced non-small-cell lung cancer with mutated EGFR: A meta-analysis from six phase III randomized controlled trials. Int J Cancer. 2012;131(5):E822–9. doi: 10.1002/ijc.27396. PubMed PMID: 22161771.

18. Antonia SJ, Villegas A, Daniel D, Vicente D, Murakami S, Hui R, Yokoi T, Chiappori A, Lee KH, de Wit M, Cho BC, Bourhaba M, Quantin X, Tokito T, Mekhail T, Planchard D, Kim YC, Karapetis CS, Hiret S, Ostoros G, Kubota K, Gray JE, Paz-Ares L, de Castro Carpeno J, Wadsworth C, Melillo G, Jiang H, Huang Y, Dennis PA, Ozguroglu M, Investigators P. Durvalumab after Chemoradiotherapy in Stage III Non-Small-Cell Lung Cancer. N Engl J Med. 2017;377(20):1919–29. doi: 10.1056/NEJMoa1709937. PubMed PMID: 28885881.

19. Gainor JF, Shaw AT, Sequist LV, Fu X, Azzoli CG, Piotrowska Z, Huynh TG, Zhao L, Fulton L, Schultz KR, Howe E, Farago AF, Sullivan RJ, Stone JR, Digumarthy S, Moran T, Hata AN, Yagi Y, Yeap BY, Engelman JA, Mino-Kenudson M. EGFR Mutations and ALK Rearrangements Are Associated with Low Response Rates to PD-1 Pathway Blockade in Non-Small Cell Lung Cancer: A Retrospective Analysis. Clin Cancer Res. 2016;22(18):4585–93. doi: 10.1158/1078-0432.CCR-15-3101. PubMed PMID: 27225694; PMCID: PMC5026567.

20. Lee CK, Man J, Lord S, Links M, Gebski V, Mok T, Yang JC. Checkpoint Inhibitors in Metastatic EGFR-Mutated Non-Small Cell Lung Cancer-A Meta-Analysis. J Thorac Oncol. 2017;12(2):403–7. doi: 10.1016/j.jtho.2016.10.007. PubMed PMID: 27765535.

21. Geng C, Paganetti H, Grassberger C. Prediction of Treatment Response for Combined Chemo- and Radiation Therapy for Non-Small Cell Lung Cancer Patients Using a Bio-Mathematical Model. Sci Rep. 2017;7(1):13542. doi: 10.1038/s41598-017-13646-z. PubMed PMID: 29051600; PMCID: PMC5648928.

22. Yagishita S, Horinouchi H, Katsui Taniyama T, Nakamichi S, Kitazono S, Mizugaki H, Kanda S, Fujiwara Y, Nokihara H, Yamamoto N, Sumi M, Shiraishi K, Kohno T, Furuta K, Tsuta K, Tamura T. Epidermal growth factor receptor mutation is associated with longer local control after definitive chemoradiotherapy in patients with stage III nonsquamous non-small-cell lung cancer. Int J Radiat Oncol Biol Phys. 2015;91(1):140–8. doi: 10.1016/j.ijrobp.2014.08.344. PubMed PMID: 25442336.

23. Edwards BK, Noone AM, Mariotto AB, Simard EP, Boscoe FP, Henley SJ, Jemal A, Cho H, Anderson RN, Kohler BA, Eheman CR, Ward EM. Annual Report to the Nation on the status of cancer, 1975-2010, featuring prevalence of comorbidity and impact on survival among persons with lung, colorectal, breast, or prostate cancer. Cancer. 2014;120(9):1290–314. doi: 10.1002/cncr.28509. PubMed PMID: 24343171; PMCID: PMC3999205.

24. Cho H, Mariotto AB, Mann BS, Klabunde CN, Feuer EJ. Assessing non-cancer-related health status of US cancer patients: other-cause survival and comorbidity prevalence. Am J Epidemiol. 2013;178(3):339–49. doi: 10.1093/aje/kws580. PubMed PMID: 23825168; PMCID: PMC3816346.

25. Enewold L, Thomas A. Real-World Patterns of EGFR Testing and Treatment with Erlotinib for Non-Small Cell Lung Cancer in the United States. PLoS One. 2016;11(6):e0156728. doi: 10.1371/journal.pone.0156728. PubMed PMID: 27294665; PMCID: PMC4905679.

26. Lim YJ, Chang JH, Kim HJ, Keam B, Kim TM, Kim DW, Paeng JC, Kang KW, Chung JK, Jeon YK, Chung DH, Wu HG. Superior Treatment Response and In-field Tumor Control in Epidermal Growth Factor Receptor-mutant Genotype of Stage III Nonsquamous Non-Small cell Lung Cancer Undergoing Definitive Concurrent Chemoradiotherapy. Clin Lung Cancer. 2017;18(3):e169–e78. doi: 10.1016/j.cllc.2016.12.013. PubMed PMID: 28131636.

27. Mak RH, Doran E, Muzikansky A, Kang J, Neal JW, Baldini EH, Choi NC, Willers H, Jackman DM, Sequist LV. Outcomes after combined modality therapy for EGFR-mutant and wild-type locally advanced NSCLC. Oncologist. 2011;16(6):886–95. doi: 10.1634/theoncologist.2011-0040. PubMed PMID: 21632451; PMCID: PMC3228219.

28. Das AK, Sato M, Story MD, Peyton M, Graves R, Redpath S, Girard L, Gazdar AF, Shay JW, Minna JD, Nirodi CS. Non-small-cell lung cancers with kinase domain mutations in the epidermal growth factor receptor are sensitive to ionizing radiation. Cancer Res. 2006;66(19):9601–8. doi: 10.1158/0008-5472.CAN-06-2627. PubMed PMID: 17018617.

29. Senan S, Brade A, Wang LH, Vansteenkiste J, Dakhil S, Biesma B, Martinez Aguillo M, Aerts J, Govindan R, Rubio-Viqueira B, Lewanski C, Gandara D, Choy H, Mok T, Hossain A, Iscoe N, Treat J, Koustenis A, San Antonio B, Chouaki N, Vokes E. PROCLAIM: Randomized Phase III Trial of Pemetrexed-Cisplatin or Etoposide-Cisplatin Plus Thoracic Radiation Therapy Followed by Consolidation Chemotherapy in Locally Advanced Nonsquamous Non-Small-Cell Lung Cancer. J Clin Oncol. 2016;34(9):953–62. doi: 10.1200/JCO.2015.64.8824. PubMed PMID: 26811519.

30. Auperin A, Le Pechoux C, Rolland E, Curran WJ, Furuse K, Fournel P, Belderbos J, Clamon G, Ulutin HC, Paulus R, Yamanaka T, Bozonnat MC, Uitterhoeve A, Wang X, Stewart L, Arriagada R, Burdett S, Pignon JP. Meta-analysis of concomitant versus sequential radiochemotherapy in locally advanced non-small-cell lung cancer. J Clin Oncol. 2010;28(13):2181–90. doi: 10.1200/JCO.2009.26.2543. PubMed PMID: 20351327.

31. Komaki R, Allen PK, Wei X, Blumenschein GR, Tang X, Lee JJ, Welsh JW, Wistuba, II, Liu DD, Hong WK. Adding Erlotinib to Chemoradiation Improves Overall Survival but Not Progression-Free Survival in Stage III Non-Small Cell Lung Cancer. Int J Radiat Oncol Biol Phys. 2015;92(2):317–24. doi: 10.1016/j.ijrobp.2015.02.005. PubMed PMID: 25968826; PMCID: PMC4432249.

32. Martinez E, Martinez M, Rico M, Hernandez B, Casas F, Vinolas N, Perez-Casas A, Domine M, Minguez J. Feasibility, tolerability, and efficacy of the concurrent addition of erlotinib to thoracic radiotherapy in locally advanced unresectable non-small-cell lung cancer: a Phase II trial. Onco Targets Ther. 2016;9:1057–66. doi: 10.2147/OTT.S89755. PubMed PMID: 27042098; PMCID: PMC4780183.

33. Ready N, Janne PA, Bogart J, Dipetrillo T, Garst J, Graziano S, Gu L, Wang X, Green MR, Vokes EE, Cancer LGBCIL. Chemoradiotherapy and gefitinib in stage III non-small cell lung cancer with epidermal growth factor receptor and KRAS mutation analysis: cancer and leukemia group B (CALEB) 30106, a CALGB-stratified phase II trial. J Thorac Oncol. 2010;5(9):1382–90. doi: 10.1097/JTO.0b013e3181eba657. PubMed PMID: 20686428.

34. Solit DB, She Y, Lobo J, Kris MG, Scher HI, Rosen N, Sirotnak FM. Pulsatile administration of the epidermal growth factor receptor inhibitor gefitinib is significantly more effective than continuous dosing for sensitizing tumors to paclitaxel. Clin Cancer Res. 2005;11(5):1983–9. doi: 10.1158/1078-0432.CCR-04-1347. PubMed PMID: 15756024.

35. Gandara DR, Gumerlock PH. Epidermal growth factor receptor tyrosine kinase inhibitors plus chemotherapy: case closed or is the jury still out? J Clin Oncol. 2005;23(25):5856–8. doi: 10.1200/JCO.2005.05.030. PubMed PMID: 16043825.

36. Warner A, Dahele M, Hu B, Palma DA, Senan S, Oberije C, Tsujino K, Moreno-Jimenez M, Kim TH, Marks LB, Rengan R, De Petris L, Ramella S, De Ruyck K, De Dios NR, Bradley JD, Rodrigues G. Factors Associated With Early Mortality in Patients Treated With Concurrent Chemoradiation Therapy for Locally Advanced Non-Small Cell Lung Cancer. Int J Radiat Oncol Biol Phys. 2016;94(3):612–20. doi: 10.1016/j.ijrobp.2015.11.030. PubMed PMID: 26867890.

37. Morgensztern D, Waqar S, Subramanian J, Gao F, Trinkaus K, Govindan R. Prognostic significance of tumor size in patients with stage III non-small-cell lung cancer: a surveillance, epidemiology, and end results (SEER) survey from 1998 to 2003. J Thorac Oncol. 2012;7(10):1479–84. doi: 10.1097/JTO.0b013e318267d032. PubMed PMID: 22982648.

38. Carmichael J, Degraff WG, Gamson J, Russo D, Gazdar AF, Levitt ML, Minna JD, Mitchell JB. Radiation sensitivity of human lung cancer cell lines. Eur J Cancer Clin Oncol. 1989;25(3):527-34. PubMed PMID: 2539297.

39. Magnuson WJ, Yeung JT, Guillod PD, Gettinger SN, Yu JB, Chiang VL. Impact of Deferring Radiation Therapy in Patients With Epidermal Growth Factor Receptor-Mutant Non-Small Cell Lung Cancer Who Develop Brain Metastases. Int J Radiat Oncol Biol Phys. 2016;95(2):673–9. doi: 10.1016/j.ijrobp.2016.01.037. PubMed PMID: 27034176.

40. Wang M, Morsbach F, Sander D, Gheorghiu L, Nanda A, Benes C, Kriegs M, Krause M, Dikomey E, Baumann M, Dahm-Daphi J, Settleman J, Willers H. EGF receptor inhibition radiosensitizes NSCLC cells by inducing senescence in cells sustaining DNA double-strand breaks. Cancer Res. 2011;71(19):6261–9. doi: 10.1158/0008-5472.CAN-11-0213. PubMed PMID: 21852385; PMCID: PMC3185115.

41. Gerlinger M, Rowan AJ, Horswell S, Math M, Larkin J, Endesfelder D, Gronroos E, Martinez P, Matthews N, Stewart A, Tarpey P, Varela I, Phillimore B, Begum S, McDonald NQ, Butler A, Jones D, Raine K, Latimer C, Santos CR, Nohadani M, Eklund AC, Spencer-Dene B, Clark G, Pickering L, Stamp G, Gore M, Szallasi Z, Downward J, Futreal PA, Swanton C. Intratumor heterogeneity and branched evolution revealed by multiregion sequencing. N Engl J Med. 2012;366(10):883–92. doi: 10.1056/NEJMoa1113205. PubMed PMID: 22397650; PMCID: PMC4878653.

42. Clevers H. The cancer stem cell: premises, promises and challenges. Nat Med. 2011;17(3):313–9. doi: 10.1038/nm.2304. PubMed PMID: 21386835.

43. Kreso A, Dick JE. Evolution of the cancer stem cell model. Cell Stem Cell. 2014;14(3):275–91. doi: 10.1016/j.stem.2014.02.006. PubMed PMID: 24607403.

44. Hata AN, Niederst MJ, Archibald HL, Gomez-Caraballo M, Siddiqui FM, Mulvey HE, Maruvka YE, Ji F, Bhang HE, Krishnamurthy Radhakrishna V, Siravegna G, Hu H, Raoof S, Lockerman E, Kalsy A, Lee D, Keating CL, Ruddy DA, Damon LJ, Crystal AS, Costa C, Piotrowska Z, Bardelli A, Iafrate AJ, Sadreyev RI, Stegmeier F, Getz G, Sequist LV, Faber AC, Engelman JA. Tumor cells can follow distinct evolutionary paths to become resistant to epidermal growth factor receptor inhibition. Nat Med. 2016;22(3):262–9. doi: 10.1038/nm.4040. PubMed PMID: 26828195; PMCID: PMC4900892.

45. Sequist LV, Waltman BA, Dias-Santagata D, Digumarthy S, Turke AB, Fidias P, Bergethon K, Shaw AT, Gettinger S, Cosper AK, Akhavanfard S, Heist RS, Temel J, Christensen JG, Wain JC, Lynch TJ, Vernovsky K, Mark EJ, Lanuti M, Iafrate AJ, Mino-Kenudson M, Engelman JA. Genotypic and histological evolution of lung cancers acquiring resistance to EGFR inhibitors. Sci Transl Med. 2011;3(75):75ra26. doi: 10.1126/scitranslmed.3002003. PubMed PMID: 21430269; PMCID: PMC3132801.

46. Engelman JA, Janne PA. Mechanisms of acquired resistance to epidermal growth factor receptor tyrosine kinase inhibitors in non-small cell lung cancer. Clin Cancer Res. 2008;14(10):2895–9. doi: 10.1158/1078-0432.CCR-07-2248. PubMed PMID: 18483355.

47. Seto T, Kato T, Nishio M, Goto K, Atagi S, Hosomi Y, Yamamoto N, Hida T, Maemondo M, Nakagawa K, Nagase S, Okamoto I, Yamanaka T, Tajima K, Harada R, Fukuoka M, Yamamoto N. Erlotinib alone or with bevacizumab as first-line therapy in patients with advanced non-squamous non-small-cell lung cancer harbouring EGFR mutations (JO25567): an open-label, randomised, multicentre, phase 2 study. Lancet Oncol. 2014;15(11):1236–44. doi: 10.1016/S1470-2045(14)70381-X. PubMed PMID: 25175099.

48. Wu YL, Lee JS, Thongprasert S, Yu CJ, Zhang L, Ladrera G, Srimuninnimit V, Sriuranpong V, Sandoval-Tan J, Zhu Y, Liao M, Zhou C, Pan H, Lee V, Chen YM, Sun Y, Margono B, Fuerte F, Chang GC, Seetalarom K, Wang J, Cheng A, Syahruddin E, Qian X, Ho J, Kurnianda J, Liu HE, Jin K, Truman M, Bara I, Mok T. Intercalated combination of chemotherapy and erlotinib for patients with advanced stage non-small-cell lung cancer (FASTACT-2): a randomised, double-blind trial. Lancet Oncol. 2013;14(8):777–86. doi: 10.1016/S1470-2045(13)70254-7. PubMed PMID: 23782814.

49. Schiller JH, Harrington D, Belani CP, Langer C, Sandler A, Krook J, Zhu J, Johnson DH, Eastern Cooperative Oncology G. Comparison of four chemotherapy regimens for advanced non-small-cell lung cancer. N Engl J Med. 2002;346(2):92–8. doi: 10.1056/NEJMoa011954. PubMed PMID: 11784875.

50. Qiao GB, Wu YL, Yang XN, Zhong WZ, Xie D, Guan XY, Fischer D, Kolberg HC, Kruger S, Stuerzbecher HW. High-level expression of Rad51 is an independent prognostic marker of survival in non-small-cell lung cancer patients. Br J Cancer. 2005;93(1):137–43. doi: 10.1038/sj.bjc.6602665. PubMed PMID: 15956972; PMCID: PMC2361489.

51. Torres-Roca JF, Eschrich S, Zhao H, Bloom G, Sung J, McCarthy S, Cantor AB, Scuto A, Li C, Zhang S, Jove R, Yeatman T. Prediction of radiation sensitivity using a gene expression classifier. Cancer Res. 2005;65(16):7169–76. doi: 10.1158/0008-5472.CAN-05-0656. PubMed PMID: 16103067.

52. Scott JG, Berglund A, Schell MJ, Mihaylov I, Fulp WJ, Yue B, Welsh E, Caudell JJ, Ahmed K, Strom TS, Mellon E, Venkat P, Johnstone P, Foekens J, Lee J, Moros E, Dalton WS, Eschrich SA, McLeod H, Harrison LB, Torres-Roca JF. A genome-based model for adjusting radiotherapy dose (GARD): a retrospective, cohort-based study. Lancet Oncol. 2017;18(2):202–11. doi: 10.1016/S1470-2045(16)30648-9. PubMed PMID: 27993569.

53. Kobe C, Scheffler M, Holstein A, Zander T, Nogova L, Lammertsma AA, Boellaard R, Neumaier B, Ullrich RT, Dietlein M, Wolf J, Kahraman D. Predictive value of early and late residual 18F-fluorodeoxyglucose and 18F-fluorothymidine uptake using different SUV measurements in patients with non-small-cell lung cancer treated with erlotinib. Eur J Nucl Med Mol Imaging. 2012;39(7):1117–27. doi: 10.1007/s00259-012-2118-8. PubMed PMID: 22526960.

54. Oka S, Uramoto H, Shimokawa H, Iwanami T, Tanaka F. The expression of Ki-67, but not proliferating cell nuclear antigen, predicts poor disease free survival in patients with adenocarcinoma of the lung. Anticancer Res. 2011;31(12):4277-82. PubMed PMID: 22199292.

55. Vesselle H, Salskov A, Turcotte E, Wiens L, Schmidt R, Jordan CD, Vallieres E, Wood DE. Relationship between non-small cell lung cancer FDG uptake at PET, tumor histology, and Ki-67 proliferation index. J Thorac Oncol. 2008;3(9):971–8. doi: 10.1097/JTO.0b013e31818307a7. PubMed PMID: 18758298.

56. Chalkidou A, Landau DB, Odell EW, Cornelius VR, O’Doherty MJ, Marsden PK. Correlation between Ki-67 immunohistochemistry and 18F-fluorothymidine uptake in patients with cancer: A systematic review and meta-analysis. Eur J Cancer. 2012;48(18):3499–513. doi: 10.1016/j.ejca.2012.05.001. PubMed PMID: 22658807.

57. Thames HD, Bentzen SM, Turesson I, Overgaard M, Van den Bogaert W. Time-dose factors in radiotherapy: a review of the human data. Radiother Oncol. 1990;19(3):219–35. doi: 10.1016/0167-8140(90)90149-q. PubMed PMID: 2281152.

58. Detterbeck FC, Gibson CJ. Turning gray: the natural history of lung cancer over time. J Thorac Oncol. 2008;3(7):781–92. doi: 10.1097/JTO.0b013e31817c9230. PubMed PMID: 18594326.

59. Switzer P, Gerstl B, Greenspoon J. Karyometry in the estimation of nuclear population in pulmonary carcinomas. J Natl Cancer Inst. 1974;52(6):1699–704. doi: 10.1093/jnci/52.6.1699. PubMed PMID: 4365386.

60. Cuaron J, Dunphy M, Rimner A. Role of FDG-PET scans in staging, response assessment, and follow-up care for non-small cell lung cancer. Front Oncol. 2012;2:208. doi: 10.3389/fonc.2012.00208. PubMed PMID: 23316478; PMCID: PMC3539654.

61. Williams MJ, Werner B, Barnes CP, Graham TA, Sottoriva A. Identification of neutral tumor evolution across cancer types. Nat Genet. 2016;48(3):238–44. doi: 10.1038/ng.3489. PubMed PMID: 26780609; PMCID: PMC4934603.

62. Bernstein L, Anderson J, Pike MC. Estimation of the proportional hazard in two-treatment-group clinical trials. Biometrics. 1981;37(3):513-9. PubMed PMID: 7317558.

63. Curran WJ, Jr., Paulus R, Langer CJ, Komaki R, Lee JS, Hauser S, Movsas B, Wasserman T, Rosenthal SA, Gore E, Machtay M, Sause W, Cox JD. Sequential vs. concurrent chemoradiation for stage III non-small cell lung cancer: randomized phase III trial RTOG 9410. J Natl Cancer Inst. 2011;103(19):1452–60. doi: 10.1093/jnci/djr325. PubMed PMID: 21903745; PMCID: PMC3186782.

64. Rohatgi A. WebPlotDigitizer V4.2. Available from: https://automeris.io/WebPlotDigitizer.

65. Foo J, Chmielecki J, Pao W, Michor F. Effects of pharmacokinetic processes and varied dosing schedules on the dynamics of acquired resistance to erlotinib in EGFR-mutant lung cancer. J Thorac Oncol. 2012;7(10):1583–93. doi: 10.1097/JTO.0b013e31826146ee. PubMed PMID: 22982659; PMCID: PMC3693219.

