## Supplemental Figures for "Modeling Resistance and Recurrence Patterns of Combined Targeted-Chemoradiotherapy Predicts Benefit of Shorter Induction Period"

### Supplementary Material:

*Full Parameter Space Analysis of Model Calibration*

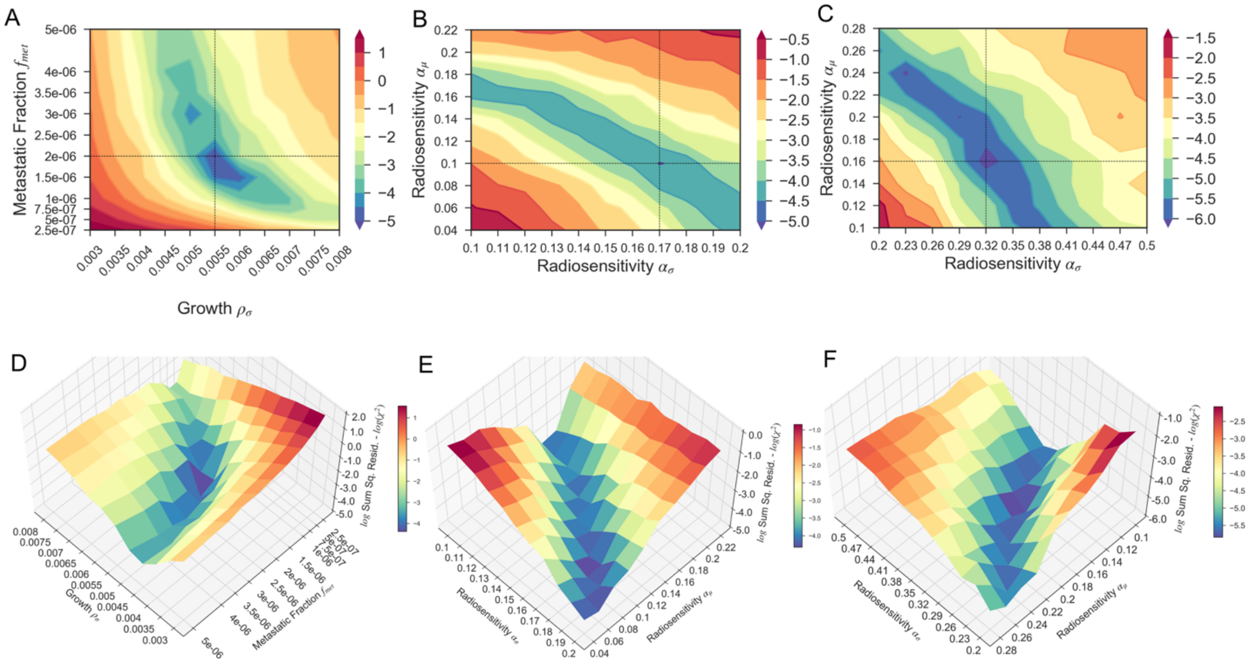


**Figure S 1:** The model parameter calibration space showing the changes in the $\log_e \chi^2$ residual as a function of initial fraction of metastatic cells and the standard deviation of the growth rate distribution (A, D), the mean and standard deviation of radiosensitivity for the wildtype (B, E) and EGFR+ (C, F) populations. In A, B, C, the parameter space is shown as a 2D contour map with the parameter values resulting in minimum $\chi^2$ residual denoted with a black dashed line. In D, E, F, the parameter space is shown as a 3D surface.

*Model Simulated PFS for WT and EGFR+ Populations Receiving Concurrent Chemoradiotherapy*

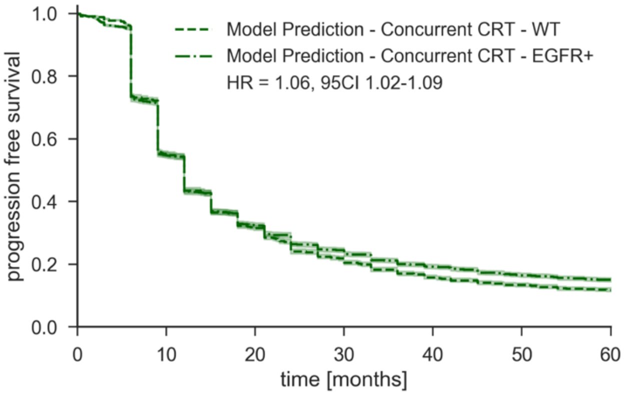


**Figure S 2:** Simulated progression free survival K-M curves for WT and EGFR+ populations with the calibrated model parameters. Even though the radiosensitivity and therefore local control in these two populations differs significantly, the similar distant failure rate leads to a similar PFS curves.

*Histograms of Model Parameters for Simulated Population*

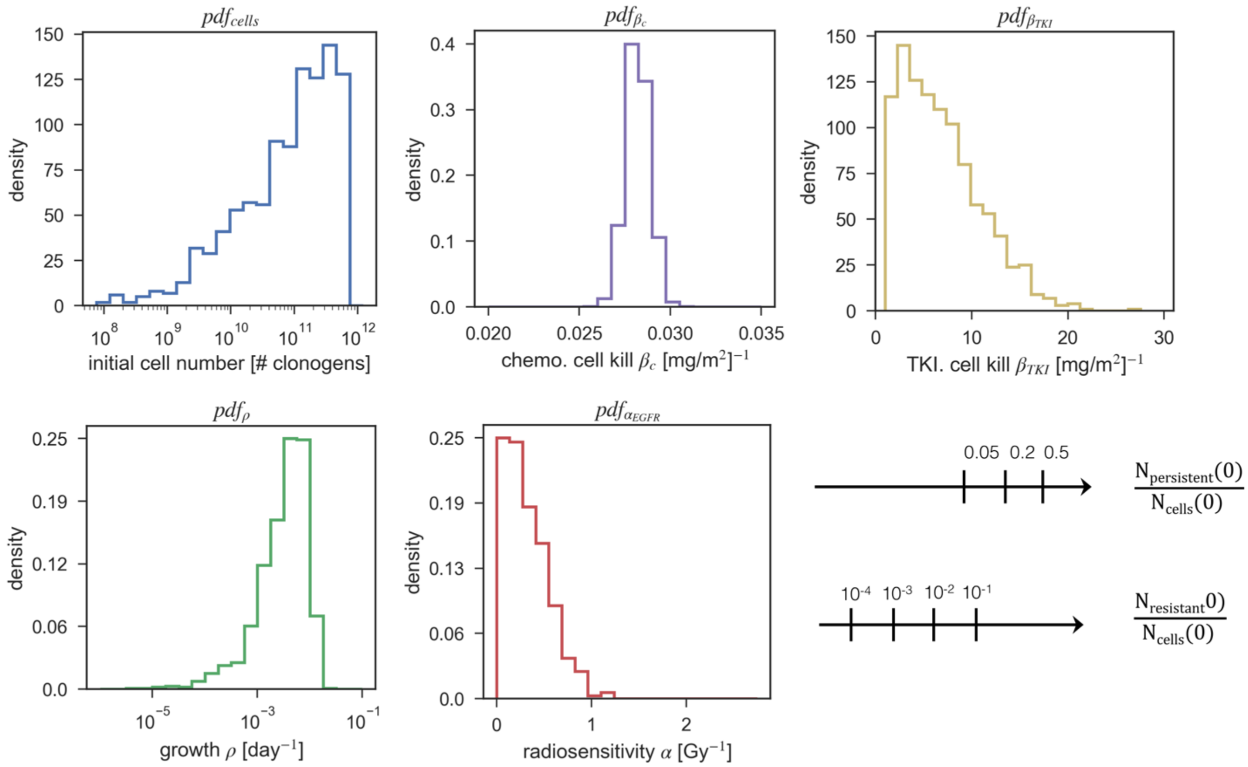


**Figure S 3:** Distributions of key model parameters used in this work for n=1024 simulated patients with 3 different initial persistent cell fractions and 4 different resistant cell fractions, yielding a total of 3x4x1024 = 12288 simulated patients.

*Model Predicted Local versus Distant Recurrence Patterns Receiving Sequential Versus Concurrent Chemotherapy*

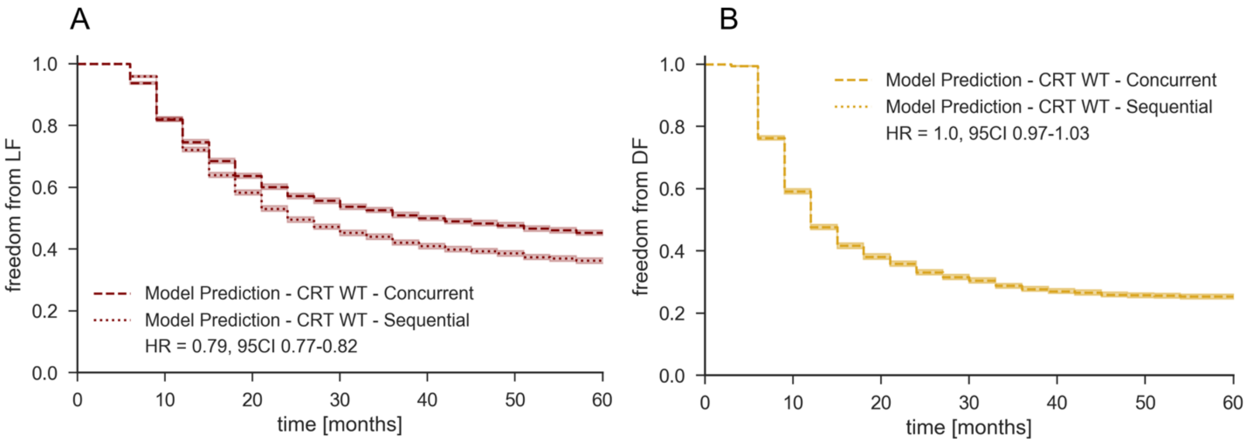


Figure S 4: Model predicted freedom from local failure (A) and distant failure (B) K-M curves for both concurrent and sequential chemoradiotherapy.

*Model Predictions of Full Multimodal Treatment Design Space*

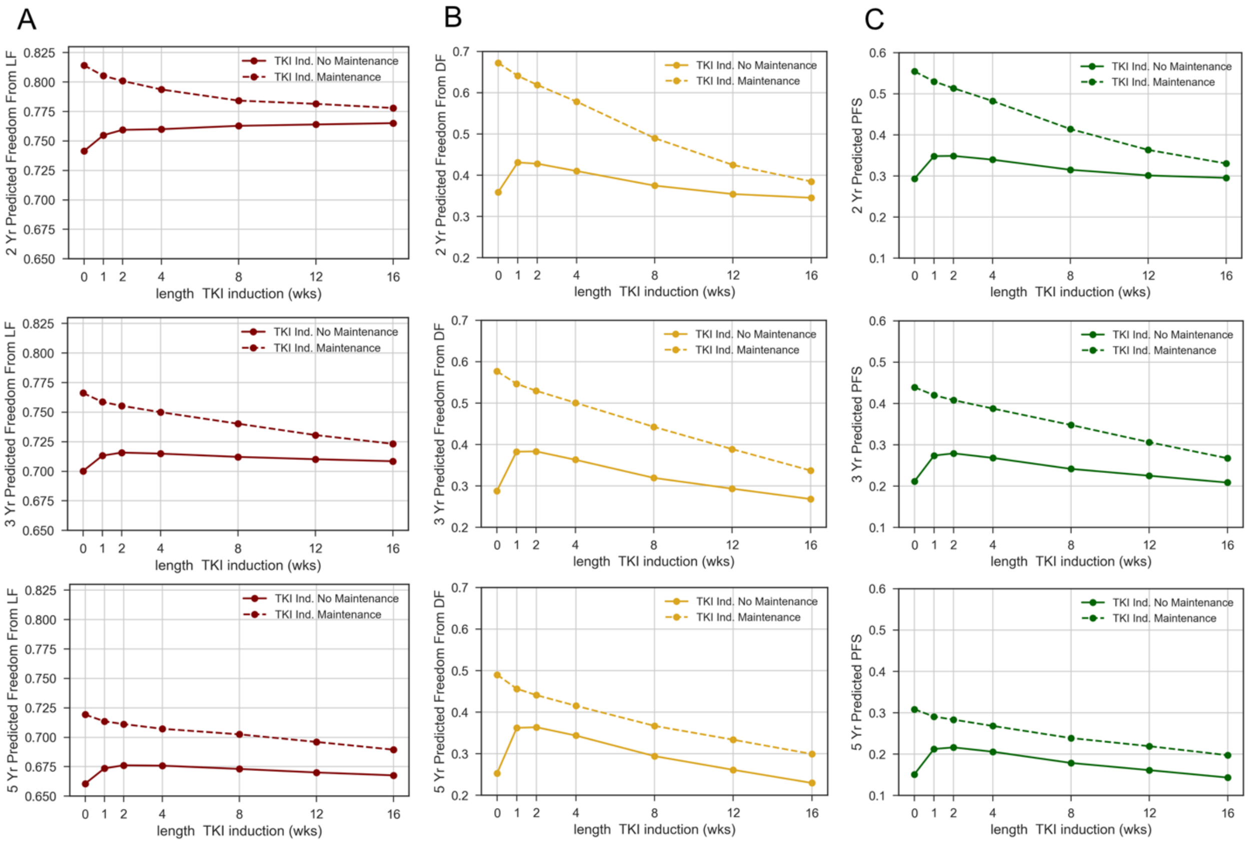


**Figure S 5:** Plots of FFLF (A), FFDF (B), and PFS (C) as function of induction length both with (solid line) and without (dashed line) adjuvant TKI maintenance at 2 yrs. (top row), 3 yrs. (middle row), and 5 yrs. (bottom row) from start of treatment.

| Group | | 2 Yr. FFLF [%] | 3 Yr. FFLF [%] | 5 Yr. FFLF [%] | Median FFLF [mo.] | 2 Yr. FFDF [%] | 3 Yr. FFDF [%] | 5 Yr. FFDF [%] | Median FFDF [mo.] | 2 Yr. PFS [%] | 3 Yr. PFS [%] | 5 Yr. PFS [%] | Median PFS [mo.] |
|---|---|---|---|---|---|---|---|---|---|---|---|---|---|
| TKI Induction + CRT | 0 wk. TKI Ind. | 74.1 | 70.0 | 66.1 | NR | 35.9 | 28.8 | 25.3 | 12.0 | 29.3 | 21.2 | 15.1 | 12.0 |
| | 1 wk. TKI Ind. | 75.5 | 71.3 | 67.4 | NR | 43.1 | 38.3 | 36.2 | 15.0 | 34.8 | 27.4 | 21.2 | 12.0 |
| | 2 wk. TKI Ind. | 75.9 | 71.6 | 67.6 | NR | 42.8 | 38.3 | 36.4 | 15.0 | 34.9 | 27.9 | 21.6 | 12.0 |
| | 4 wk. TKI Ind. | 76.0 | 71.5 | 67.6 | NR | 41.0 | 36.3 | 34.4 | 15.0 | 34.0 | 26.8 | 20.6 | 12.0 |
| | 8 wk. TKI Ind. | 76.3 | 71.2 | 67.3 | NR | 37.5 | 31.9 | 29.4 | 15.0 | 31.5 | 24.2 | 17.9 | 12.0 |
| | 12 wk. TKI Ind. | 76.4 | 71.0 | 67.0 | NR | 35.4 | 29.3 | 26.1 | 15.0 | 30.1 | 22.5 | 16.1 | 15.0 |
| | 16 wk. TKI Ind. | 76.5 | 70.8 | 66.8 | NR | 34.5 | 26.8 | 23.0 | 15.0 | 29.5 | 20.9 | 14.3 | 15.0 |
| TKI Induction + CRT + TKI Maintenance | 0 wk. TKI Ind. | 81.4 | 76.6 | 71.9 | NR | 67.2 | 57.7 | 49.0 | 54.0 | 55.5 | 43.9 | 30.9 | 27.0 |
| | 1 wk. TKI Ind. | 80.5 | 75.9 | 71.4 | NR | 64.1 | 54.7 | 45.6 | 42.0 | 53.0 | 42.0 | 29.1 | 25.9 |
| | 2 wk. TKI Ind. | 80.1 | 75.5 | 71.1 | NR | 61.9 | 53.0 | 44.1 | 39.0 | 51.4 | 40.8 | 28.3 | 24.0 |
| | 4 wk. TKI Ind. | 79.4 | 75.0 | 70.7 | NR | 57.9 | 50.1 | 41.5 | 36.0 | 48.3 | 38.8 | 26.8 | 21.0 |
| | 8 wk. TKI Ind. | 78.4 | 74.0 | 70.2 | NR | 49.0 | 44.3 | 36.7 | 21.0 | 41.4 | 34.8 | 23.9 | 15.0 |
| | 12 wk. TKI Ind. | 78.1 | 73.1 | 69.6 | NR | 42.5 | 38.9 | 33.4 | 15.0 | 36.3 | 30.6 | 21.9 | 15.0 |
| | 16 wk. TKI Ind. | 77.8 | 72.3 | 68.9 | NR | 38.5 | 33.7 | 29.9 | 15.0 | 33.0 | 26.7 | 19.7 | 15.0 |

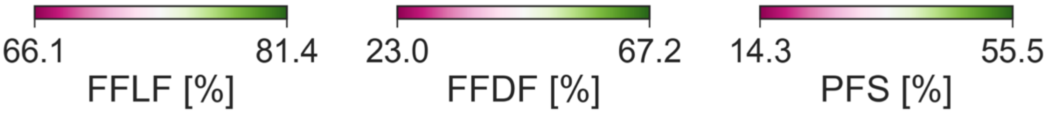


**Figure S 6**: Table of outcomes for complete multimodal treatment design parameter space, color coded by range of effect. NR= not reached.

*Estimated Power of Varying Induction Length on PFS as a Function of Sample Size*

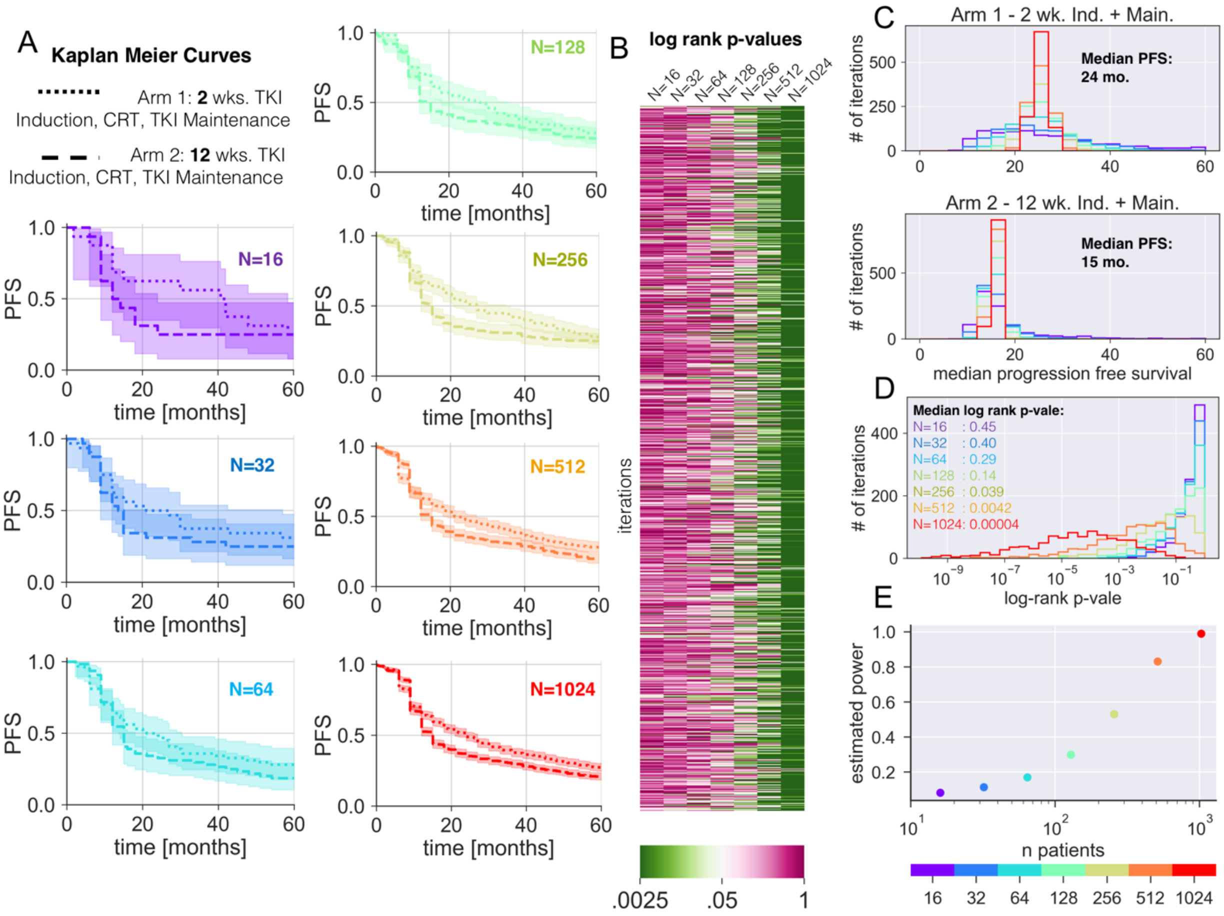


**Figure S 7:** Simulated PFS K-M curves for 2 wk. and 12 wk. induction lengths with increasing number of simulated patients. A heatmap of log rank p-values testing statistical difference between the 2 wk. versus 12 wk. PFS K-M curves for 1000 iterations of the simulation at each sample size. Histograms of the median PFS (C) and log rank p-values (D) for the 1000 iterations of the 2 wk. and 12 wk. induction simulations at each sample size. Estimated statistical power as a function of sample size. Here statistical was estimated as the fraction of iteration resulting in a p-value<0.05.
